# A Blast Fungus Effector Targets a Tubby-like F-box Protein to Suppress Plant Immunity

**DOI:** 10.64898/2026.10.15.759101

**Authors:** Indira Saado, Josh W Bennett, Luyao Huang, Khong-Sam Chia, Nicholas J Talbot, Mark J Banfield

**Affiliations:** Department of Biochemistry and Metabolism, John Innes Centre, Norwich Research Park, Norwich, NR4 7UH, UK; The Sainsbury Laboratory, University of East Anglia, Norwich Research Park, Norwich, NR4 7UH, UK Norwich, UK; Department of Molecular Biology, Max Planck Institute for Biology Tübingen, 72076 Tübingen, Germany

## Abstract

The blast fungus *Magnaporthe oryzae* is one of the most destructive pathogens of rice, secreting effector proteins that reprogram host cells to enable infection. Identifying plant targets of these fungal effectors is central to understanding how blast disease is established, but the targets and mechanisms of most effectors remain unknown. Here, we show that a Zinc finger (ZiF)-family effector, Tie1, targets the rice Tubby-like protein (TLP) family, which are membrane-associated plant immunity regulators whose exact functions have not been defined. Tie1 binds a range of rice TLPs and specifically engages with the Tubby-like domain of OsTLP7. Co-expression *in planta* with Tie1 re-localises this domain from the plasma membrane to the cytoplasm and nucleus, a shift also observed by disrupting the OsTLP7 phosphatidylinositol 4,5-bisphosphate (PIP_2_)-binding site in the absence of the effector. By combining AlphaFold3 structural predictions with allelic variation, we define a putative Tie1/OsTLP7 interface distal to the PIP_2_-binding site. Furthermore, Tie1 induces transcriptional activation activity of the OsTLP7 Tubby-like domain, and Tie1 overexpression in stable *Arabidopsis* lines leads to upregulation of genes that repress jasmonic acid (JA) signalling. Taken together, our results indicate that Tie1 inhibits PIP_2_ binding by OsTLP7 to drive its re-localisation to the nucleus and thereby modulating transcription to suppress JA-mediated plant immunity.

**SIGNIFICANCE STATEMENT:** Plant pathogens secrete effector proteins that reprogram host cells to promote disease. Understanding how effectors contribute to virulence is therefore critical to developing new forms of disease resistance. Tubby-like proteins (TLPs) are conserved membrane-associated proteins with roles in plant immunity that are poorly understood. Here we identify an effector from the devastating rice blast fungus, Tie1, that targets rice TLPs disrupting membrane localisation of OsTLP7, sending the protein to the nucleus where it alters transcriptional activity. We show that Tie1 upregulates genes that suppress jasmonic acid-mediated plant immunity. This work reveals how TLPs are targeted by pathogens to suppress immunity.

## INTRODUCTION

*Magnaporthe oryzae* is a destructive fungal plant pathogen that deploys a large repertoire of more than 500 secreted proteins called effectors to manipulate host immunity and promote infection in cereal crops including rice, wheat and barley (1). Typically, effectors display high levels of positive selection, resulting in functional diversification and evasion of detection by the plant immune system (2). However, despite low sequence similarity, it is now well established that many plant pathogen effectors cluster into structurally related families (3–8). These families may arise through gene duplication followed by selection pressure generating sequence diversification while conserving the core effector fold (8, 9). This evolutionary process can lead to the functional diversification of effectors, including the manipulation of novel host targets to promote infection. However, the molecular mechanisms by which most sequence-unrelated structurally similar (SUSS) *Magnaporthe* effectors contribute to virulence and disease development on infection remains poorly understood (10).

Genetic mapping of recognition by intracellular plant immune receptors, including nucleotide binding leucine-rich repeat (NLR) proteins and kinase-fusion proteins (KFPs), is a common mechanism to discover effectors. This has been a powerful approach for cloning *Magnaporthe* effectors, for example, AVR-Pik, AVR-Pia, AVR1-CO39, AVR-Mgk1, AVR-Piz-t, AVR-Pib, PBY2, PWT3 and PWT7, all of which belong to the *Magnaporthe* Avrs and ToxB (MAX) fold, and AVR-Pii, a zinc-finger fold (ZiF) effector (11–17). Many other *Magnaporthe* effectors are predicted to adopt MAX and ZiF folds but have not had a corresponding immune receptor identified. Most MAX effector/immune receptor interactions identified are mediated via integrated heavy metal-associated (HMA) domains in the receptors, which also suggests plant HMA domain-containing proteins are putative virulence-associated host targets (18–21). The interaction of the ZiF effector AVR-Pii with rice Exo70F2/F3 is required for immune perception and highlights Exo70 as a virulence target for this effector (22, 23). However, virulence-associated targets for hundreds of other predicted MAX and ZiF effectors are unknown, and this knowledge is required to understand how *Magnaporthe* effectors manipulate host cells to cause disease.

The SUSS ZiF effector family has been classified into ten tribes based on protein sequence, with AVR-Pii belonging to Tribe VIII (24). Several ZiF effectors belonging to Tribe IV have been shown to modulate transcription of host defence-associated genes by directly binding to DNA (25–27). Other diverse effectors across different pathogens also manipulate host transcription (e.g. Nkd1, TALEs, and CRNs (28–30)), underlining transcriptional reprogramming as a strategy employed by plant pathogens to colonise their hosts (31). This includes modifying the function of transcription factors as well as direct DNA binding. However, to date, effector interaction with a conserved yet understudied group of membrane bound plant stress-related regulators - the Tubby-like F-box proteins (TLPs) - has not been demonstrated. TLPs are proposed to function as intermediates connecting plasma membrane signalling to nuclear transcriptional regulation through a phosphatidylinositol 4,5-bisphosphate (PIP₂)-binding mechanism. Upon activation, they can be released from the membrane and translocate to the nucleus, where they modulate transcriptional responses (32). In plants, TLPs have been implicated in diverse biological processes, including growth and development, abiotic and biotic stress responses, and hormone signalling (32, 33). Although conserved across eukaryotic species, TLP involvement in plant biotic stress responses remains underexplored, however, some examples have been documented. For example, expression of all 14 TLPs in rice is upregulated in response to infection by the bacterial pathogen *Xanthomonas oryzae pv. oryzae* (34). Further, OsTLP2 regulates expression of *OsWRKY13* by binding to a *cis*-element PRE4 (35). *OsWRKY13* has been implicated in pathogen resistance as transgenic rice plants overexpressing this gene exhibit enhanced resistance to blast disease (36). TLPs also have a demonstrated role in proteolytic regulation in wheat, TaTLP5 is involved in degradation of the susceptibility factor Catalase1 (which decreases ROS production) to reinforce resistance to stripe rust (37). Collectively, characterised TLPs have diverse functions with emerging roles in regulating immune signalling.

In this study, we establish the interaction of a ZiF family Tribe I effector with the host Tubby-like F-box protein family. We show that the <u>T</u>ubby-like F-box protein interacting <u>e</u>ffector <u>1</u> (Tie1) from *M. oryzae* binds to a range of rice TLPs (OsTLPs), specifically targeting the Tubby-like domain of OsTLP7. Co-expression of Tie1 and OsTLP7 in *Nicotiana benthamiana* leads to relocalisation of the Tubby-like domain of OsTLP7 (OsTLP7-TLD) from the plasma membrane to the cytoplasm and nucleus. A similar localisation pattern is also observed upon mutation of the OsTLP7 PIP_2_ binding site in the absence of the effector. Leveraging AlphaFold3 structural predictions and allelic variation, we define a putative Tie1/OsTLP7 binding interface, distal to the PIP_2_ binding site. Furthermore, we demonstrate through GAL4-tagging that Tie1 interferes with transcriptional activation activity mediated by the OsTLP7-TLD in *N. benthamiana*. Finally, in stable *Arabidopsis* lines overexpressing Tie1, we observe upregulation of genes involved in repressing Jasmonic acid signalling. Together, these results provide evidence that Tie1 acts as an inhibitor of PIP_2_ binding of the OsTLP7-TLD to promote relocalisation from the plasma membrane to the nucleus to induce transcriptional reprogramming. These data represent a new virulence strategy deployed by the rice blast fungus during infection.

## RESULTS

### Tie1 is a BIC-localised effector

Tie1 (previously named ZiFIj) is a *M. oryzae* effector that belongs to Tribe I of the ZiF family (24). The wild type rice pathogenic *M. oryzae* strain, Guy11, contains two closely related *Tie1* homologues, designated *Tie1k* and *Tie1f*. Published transcriptome data showed that both homologs are expressed during infection, although *Tie1f* accumulated at markedly higher levels during the biotrophic phase (38) (Fig. S1A). To examine Tie1f localisation during infection, a Tie1f-GFP fusion driven by its native promoter was introduced into *M. oryzae*. During rice leaf sheath infection, Tie1f-GFP fluorescence was detected in invasive hyphae and accumulated at the biotrophic interfacial complex (BIC) (Fig. S1B), a specialised structure associated with the delivery of cytoplasmic effectors into host cells. Co-localisation analyses with the fluorescently tagged cytoplasmic effector marker Pwl2 and apoplastic effector marker Slp1 showed that Tie1f-GFP co-localised with Pwl2 but not with Slp1 (Fig. 1A), indicating that Tie1f behaves as a cytoplasmic effector that accumulates at the BIC during infection.

**Figure 1.**
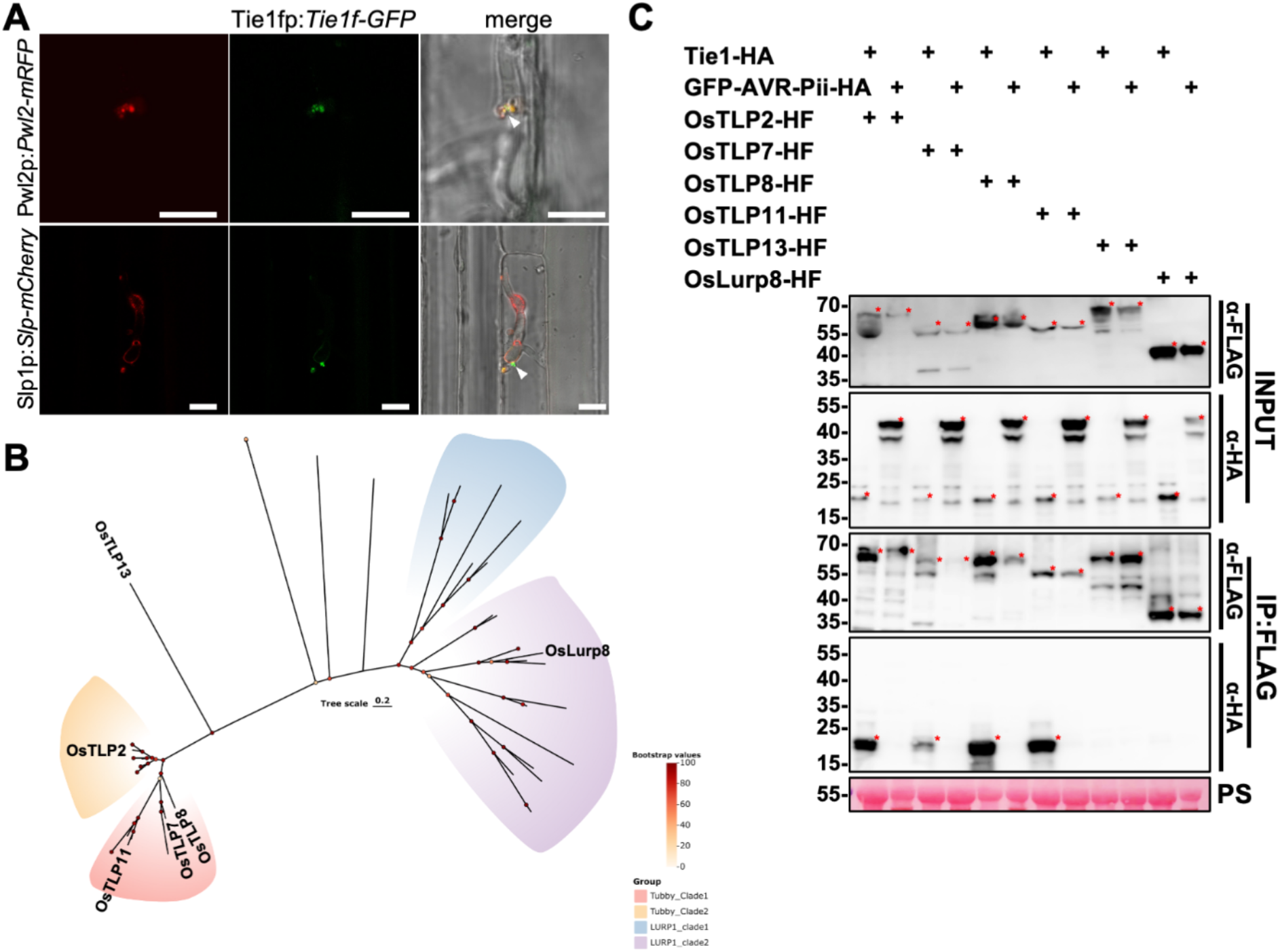
Tie1 is a *M. oryzae* BIC-localised effector that interacts with diverse plant Tubby-like proteins. (A) Localisation of Tie1f-GFP during rice leaf sheath infection. Tie1f-GFP accumulated at the biotrophic interfacial complex (BIC) and co-localised with the cytoplasmic effector marker Pwl2 but not with the apoplastic effector marker Slp1. Images were acquired at 26–28 h post-inoculation. Scale bars, 20 μm. (B) Phylogenetic analysis of the rice Tubby-like protein (TLP) family. TLP proteins selected for interaction analyses are indicated. Branch colours indicate bootstrap values ranging from 0 (yellow) to 100 (dark red). (C) Co-immunoprecipitation assays between Tie1-HA and representative rice Tubby-like proteins expressed in *Nicotiana benthamiana*. OsTLP2-HF, OsTLP7-HF, OsTLP8-HF, OsTLP11-HF, OsTLP13-HF, and LURP8-HF were immunoprecipitated using anti-FLAG resin, and associated Tie1-HA was detected by immunoblotting. Interactions were detected with OsTLP2, OsTLP7, OsTLP8, and OsTLP11, but not with OsTLP13 or LURP8. GFP-AVR-Pii-HA was included as a specificity control and showed no interaction with the tested Tubby-like proteins. Red asterisks indicate the bands of appropriate molecular mass for the expressed protein. Ponceau S staining indicates total protein loading in the input samples. The experiment was repeated three times with similar results.

### Tie1 interacts with diverse plant Tubby-like proteins

To determine potential host targets of Tie1 we used a co-immunoprecipitation/mass spectrometry (IP/MS) approach. For this, Tie1-GFP (without the predicted signal peptide, comprising residues 1-19) was transiently expressed in *N. benthamiana* leaves. Leaf tissue transiently expressing GFP alone was used as a negative control. Subsequent mass spectrophotometry (IP/MS) analysis identified Tubby-like F-box proteins as strongly enriched in the Tie1-GFP samples compared to GFP alone, repeatedly ranking as top candidate interactors (Fig. S2, Table S1). The repeated recovery of *N. benthamiana* TLP proteins upon expression with Tie1 suggested that members of this family may represent conserved host targets of this effector.

To link this finding to a host plant of *Magnaporthe oryzae*, we explored the diversity of Tubby-like domain-containing proteins in rice by constructing a phylogenetic tree (Fig. 1B). This analysis revealed two distinct clades: LURPs (*LATE UPREGULATED IN RESPONSE TO HYALOPERONOSPORA PARASITICA*), which harbour a canonical Tubby-like fold (39, 40), and TLPs, that possess a Tubby-like fold in combination with an N-terminal F-box domain (41, 42) (Fig. S3A, B). Representative members across the phylogeny with various domain architectures were selected for interaction assays. Co-immunoprecipitation experiments (using AVR-Pii as a negative control) in *N. benthamiana* showed that Tie1 interacts with OsTLP2, OsTLP7, OsTLP8, and OsTLP11, whereas no interaction was detected with OsTLP13 or LURP8 (Fig. 1C). Sequence comparison identified OsTLP7 as the closest rice homologue to the most prevalent TLP hit from the IP/MS (NaTLP3) with OsTLP7 subsequently used to assess the specificity of the TLP/Tie1 interaction (Fig. S4A). Co-IP analysis confirmed that Tie1 does not associate with previously characterised effector targets such as Exo70F3 or HIPP43, and that OsTLP7 does not interact with other unrelated *M. oryzae* effectors (Fig. S4B). These findings consequently led us to assess the contribution of Tie1 to pathogenicity. Single and double deletion mutants of the two Guy11 Tie1 homologues were generated using CRISPR–Cas9-mediated genome editing (Fig. S5A). Spray inoculation assays on the susceptible rice cultivar CO-39 revealed no detectable differences in lesion development between mutant and wild-type strains, as reported for many effector-encoding genes and likely a consequence of extensive redundancy in the effector repertoire (Fig. S5B). We conclude that OsTLP7 constitutes a host target for the *M. oryzae* ZiF family effector Tie1.

### Tie1 specifically interacts with and re-localises the Tubby-like domain of OsTLP7

To determine whether a specific domain of OsTLP7 mediates interaction with Tie1, truncations of OsTLP7 were designed corresponding to the F-box domain (OsTLP7-FboxD, residues 1-100) and the Tubby-like domain (OsTLP7-TLD, residues 101-406) (Fig. S3). Co-IP assays were performed using the truncated constructs, together with Tie1, Pwl2, or AVR-Pii. Full-length OsTLP7 co-expressed with Tie1 served as a positive control. Tie1, but not the other effectors, co-immunoprecipitated with the Tubby-like domain, whereas no interaction was detected with OsTLP7-FboxD for any effector (Fig. 2A). This demonstrates that Tie1 specifically interacts with the Tubby-like domain of OsTLP7.

**Figure 2:**
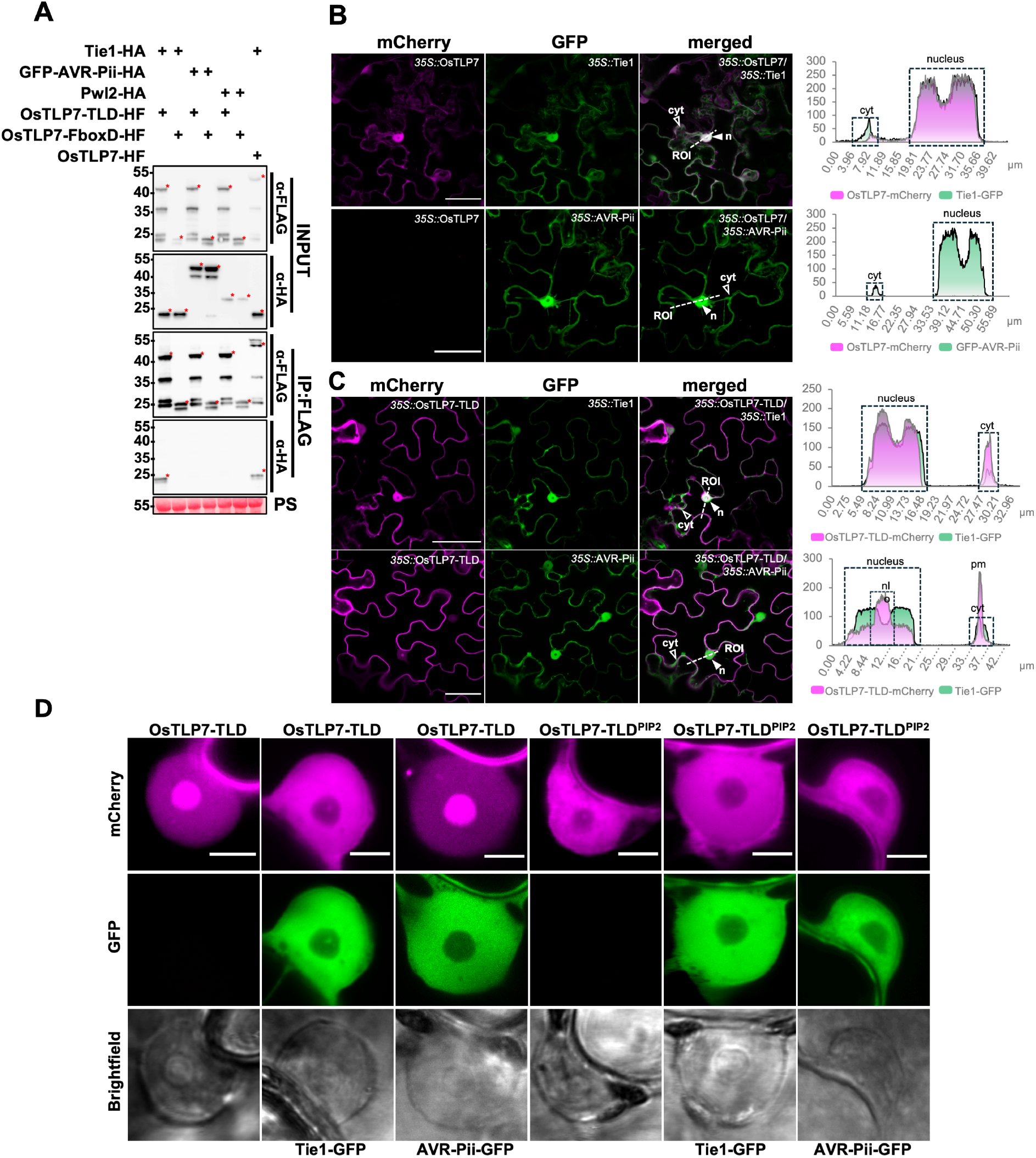
Tie1 binds the OsTLP7 Tubby domain and alters its localisation in plant cells. (A) Co-immunoprecipitation assays were performed using individual domains of OsTLP7-HF, the F-box domain, and the Tubby domain. Tie1-HA specifically interacted with the Tubby domain, while no binding was observed with the F-box domain. No interactions were detected with GFP-AVR-Pii-HA or Pwl2-HA. Red asterisks indicate the bands of appropriate molecular mass for the expressed protein. Ponceau S staining was used to confirm equal protein loading in the input samples. The experiment was repeated three times with consistent results. (B) Representative images of OsTLP7-mCherry co-localised with either Tie1-GFP or GFP-AVR-Pii (control). OsTLP7 co-expressed with Tie1 is detectable in both the cytoplasm and nuclei (n, filled white arrowhead highlights representative nucleus; cyt, blank arrowhead highlights representative cytoplasmic regions). OsTLP7 co-expressed with AVR-Pii shows no detectable signal. Tie1 and AVR-Pii both localise to the cytoplasm and nuclei. Region of interest (ROI, merged images) indicate starting points of transection lines used for fluorescence intensity profiling. Right panels: Fluorescence intensity plots along transection lines for GFP and mCherry channels. Nuclear areas and peripheral cytoplasmic signals are boxed. Maximum fluorescence intensity = 255. *Note: Only relative presence and spatial changes in intensity can be compared between fluorophores; overall signal intensities may differ.* Scale bar = 50 µm. (C) Representative images of OsTLP7-TLD-mCherry co-localised with either Tie1-GFP or GFP-AVR-Pii (control). OsTLP7-TLD with AVR-Pii is mainly localised at the plasma membrane and nucleus. Tie1 and AVR-Pii both localise to the cytoplasm and nuclei. Region of interest (ROI, merged images) indicate starting points of transection lines used for fluorescence intensity profiling. Right panels: Fluorescence intensity plots along transection lines for GFP and mCherry channels. Nuclear areas and peripheral cytoplasmic signals are boxed. Maximum fluorescence intensity = 255. *Note: Only relative presence and spatial changes in intensity can be compared between fluorophores; overall signal intensities may differ.* Scale bar = 50 µm. (D) Representative localisation patterns of OsTLP7-TLD and OsTLP7-TLD^PIP2^ in single plant nuclei, expressed alone or co-expressed with Tie1 or AVR-Pii. OsTLP7-TLD alone, and OsTLP7-TLD co-expressed with AVR-Pii, predominantly localise to the nucleolus, whereas OsTLP7-TLD co-expressed with Tie1 localises throughout the nucleus. In contrast, OsTLP7-TLD^PIP2^ shows a consistent distribution in plant nuclei, regardless of expressing it alone or co-expression with Tie1 or AVR-Pii. Scale bars = 5 µm.

As the localisation of Tubby domains is known to influence their function, we next examined the subcellular localisation of OsTLP7, and how this is influenced by Tie1, in plant cells. When *N. benthamiana* leaves expressing OsTLP7-mCherry were analysed by confocal microscopy, no detectable mCherry fluorescence was observed yet co-expressed NLS-CFP (nuclear localisation signal) fluorescence could be perceived in nuclei, confirming that analysed cells were viable (Fig. 2B, S6A). Intriguingly, when OsTLP7-mCherry was co-expressed with Tie1-GFP, a strong mCherry signal was detected, with OsTLP7 co-localising with Tie1 in both the cytoplasm and nucleus throughout transformed tissue (Fig. 2B). Co-expression of OsTLP7-mCherry with the non-interacting effector GFP-AVR-Pii showed the same result as OsTLP7-mCherry alone, with no accumulation of OsTLP7-mCherry being observed in microscopy or western blot (Fig. 2B, S6B). Taken together, these results suggest that Tie1 interacts with and stabilises OsTLP7 in plant cells.

To investigate the impact of Tie1 interaction on the individual domains of OsTLP7, OsTLP7-TLD and OsTLP7-FboxD were each fused to mCherry (at the C-terminus) and co-expressed with Tie1 or AVR-Pii. Consistent with previous literature on Tubby-domains, OsTLP7-TLD-mCherry predominantly localised to the plasma membrane, overlapping with plasma membrane markers (33, 43) (Fig. 2C, S6C). By contrast, when OsTLP7-TLD-mCherry was co-expressed with Tie1-GFP, it accumulated in the cytoplasm (Fig. 2C). Higher magnification of the nucleus further revealed that OsTLP7-TLD-mCherry accumulated in the nucleolus in the presence of GFP-AVR-Pii, whereas co-expression with Tie1-GFP resulted in accumulation in the nucleoplasm (Fig. 2D). By contrast, OsTLP7-FboxD-mCherry maintained a diffuse localisation pattern in plant cells, without notable changes when co-expressed with either GFP-AVR-Pii or Tie1-GFP (Fig. S6B). Accumulation of all proteins was verified by immunoblotting (Fig. S6D).

Tubby-like domains are frequently anchored to the plasma membrane via binding to the phospholipid, phosphatidylinositol 4,5-bisphosphate (PIP₂) (44). The crystal structure of a mouse Tubby protein bound to PIP_2_ (PDB: 1I7E) identified two residues vital for binding to the phospholipid (44), which are conserved in OsTLP7-TLD (Lys189 and Arg191). To investigate the importance of these residues in OsTLP7-TLD, both were mutated to alanine residues (Lys189Ala and Arg191Ala, OsTLP7-TLD^PIP2^). Following expression, we observed the subcellular localisation of OsTLP7-TLD^PIP2^-mCherry was predominantly in the cytoplasm and nucleoplasm, and was unaffected by co-expression with either AVR-Pii or Tie1 (Fig. 2D, S7A-B). This confirmed that the mutated residues are critical for plasma membrane association in OsTLP7-TLD. We hypothesised that disrupting the PIP₂-binding interface in OsTLP7-TLD could affect its interaction with Tie1, given that Tie1 induces re-localisation of this domain from the plasma membrane on co-expression. However, OsTLP7-TLD^PIP2^ retained association with Tie1 following pull-down in co-IP assays, similar to wild-type (Fig. S7C). These experiments suggest that the Tie1 effector does not prevent OsTLP7-TLD binding to PIP_2_ through direct interaction with these motif residues, and likely induces re-localisation through another mechanism.

### Tie1/OsTLP7 interaction can be disrupted based on AlphaFold3 predictions and effector sequence diversity

After establishing that Tie1 specifically binds to OsTLP7-TLD, but most likely does not bind to the OsTLP7 PIP_2_-binding site, we sought to locate the binding interface between these proteins. Firstly, we used AlphaFold3 to predict the complex of OsTLP7-TLD with Tie1. For this we used only the zinc finger fold region of Tie1 (Tie1^153^, which we confirmed retained binding to OsTLP7-TLD (Fig. S8A-B)). The interaction of OsTLP7-TLD with Tie1^153^ was modelled across three independent runs obtaining ipTM scores of 0.6, 0.68 and 0.54 (Fig. S8C and S9A). The predictions consistently identified the same interacting residues for the OsTLP7-TLD, positioned on an extended anti-parallel β-sheet structure protruding from the core β-barrel (Fig. 3A). By contrast, the orientation of Tie1 at this site was not consistently predicted across models (Fig. S9A). Despite this, based on the most confident prediction, we generated mutants for both OsTLP7-TLD (Cys331Glu, Phe336Glu, Val340Glu, and As347Glu) and Tie1 (Tyr56Arg, Val58Glu, Asn63Arg, Gln65Arg) and probed for interaction by co-IP (Fig. 3B and S9C). We observed that no Tie1 mutant was sufficient to disrupt binding to OsTLP7-TLD (Fig. S9C). However, all four single OsTLP7-TLD mutants abolished interaction with Tie1 (Fig. 3B). Intriguingly, unlike the native protein, the non-interacting mutant OsTLP7-TLD^F336E^ mutant localised to the cytoplasm and nucleolus rather than the plasma membrane, while Tie1 failed to re-localise it to the nucleus (Fig. S10A-B). This suggests that the extended anti-parallel β-sheet region of OsTLP7, identified as an interaction interface by AlphaFold3, is involved in localisation of the wild-type protein to the plasma membrane and its interaction with Tie1.

**Figure 3:**
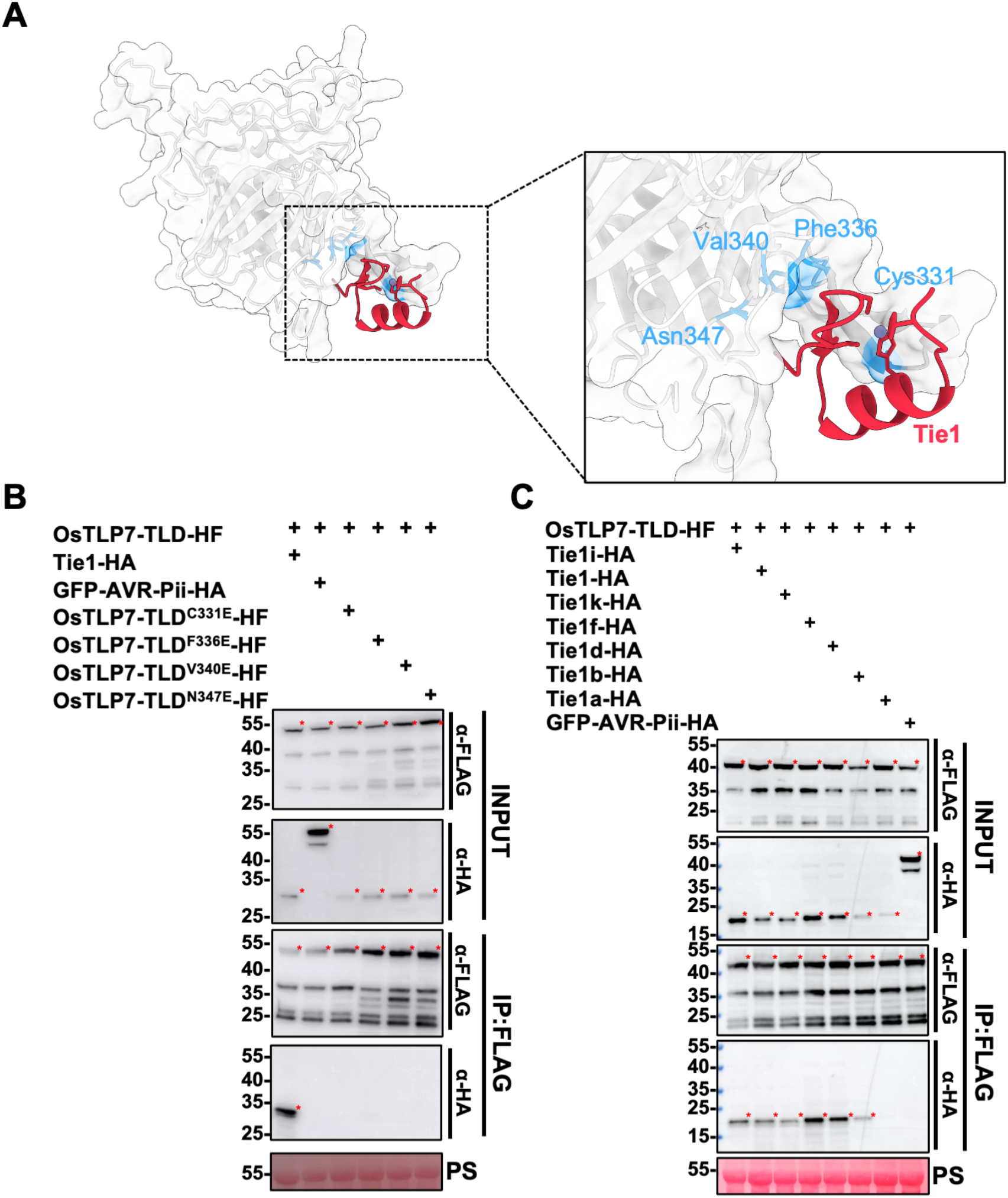
AlphaFold3 prediction-based mutation combined with allelic diversity identifies an interaction interface. (A) AlphaFold3 model of the predicted Tie1 (red):OsTLP7-TLD (gray) interface with residues of the OsTLP7-TLD (blue) predicted to interact with Tie1 highlighted. (B) Co-immunoprecipitation assays of four mutant OsTLP7-TLD-HF proteins with Tie1, demonstrating loss of binding to all OsTLP7-TLD mutants. No interactions were detected between OsTLP7-TLD-HF with GFP-AVR-Pii-HA. Red asterisks indicate the bands of appropriate molecular mass for the expressed protein. Ponceau S staining was used to confirm equal protein loading in the input samples. The experiment was repeated three times with consistent results. (C) Co-immunoprecipitation assays of OsTLP7-TLD-HF with selected Tie1-HA clade effectors, demonstrating loss of binding only for the Tie1a allele; all other alleles retain binding to OsTLP7-TLD. No interactions were detected between OsTLP7-TLD-HF and GFP-AVR-Pii-HA. Red asterisks indicate the bands of appropriate molecular mass for the expressed protein. Ponceau S staining was used to confirm equal protein loading in the input samples. The experiment was repeated three times with consistent results.

As AlphaFold3 was unable to predict the interacting residues on Tie1, we explored the Tie1 effector tribe as a source of diversity for OsTLP7-TLD binding. An unrooted phylogeny of Tie1 tribe members (previously annotated as Tribe I (24)) was generated (Fig. S11A) and a representative subset of effectors spanning phylogenetic diversity was selected to assess interactions with the OsTLP7-TLD by co-IP. Interestingly, all tested Tie1 tribe effectors bound to OsTLP7-TLD, with the exception of Tie1a (Fig. 3C). To identify determinants underlying this differential interaction, the amino acid sequences of Tie1 and Tie1a were compared and four Tie1/Tie1a chimeras were generated (Fig. S11B). The ability of these chimeras to bind OsTLP7-TLD tested using a co-IP approach. These assays showed that two chimeras, sharing common substitutions His60Arg and Ala61Asp (HA/RD) (chimera 1 and 3), lost binding to OsTLP7-TLD (Fig. S11C). To test the impact of these mutants directly, His60Arg and Ala61Asp substitutions were introduced into Tie1 together (Tie1^RD^), and the interaction with OsTLP7-TLD probed by co-IP. The Tie1^RD^ variant consistently displayed reduced band strength upon pull-down with OsTLP7-TLD, compared to wild-type Tie1, but did not completely abolish binding (Fig. S11D). The HA/RD residues are located between two cysteine residues (Cys59 and Cys62) that coordinate the Zn^2+^ ion; therefore, we investigated whether this may disrupt the zinc finger fold. To test this, we generated a triple CCH mutant of Tie1 (Cys59Ala, Cys62Ala and His75Ala) to abolish Zn^2+^ binding, which leads to protein instability in AVR-Pii (24, 45). While Tie1^RD^ protein accumulated to similar levels as wildtype Tie1, we did not observe accumulation of Tie1^CCH^ (Fig. S11D). This suggests that the His60Arg and Ala61Asp mutations in Tie1 do not lead to protein instability through disrupting zinc coordination and may directly contribute to binding OsTLP7-TLD.

### Tie1 induces OsTLP7-TLD transcriptional activation activity and overexpression of Tie1 upregulates suppressors of Jasmonic acid signalling

Tubby domain-containing proteins function as transcriptional activators, and therefore tagging these proteins with the GAL4 DNA-binding domain should activate expression of a reporter gene downstream of the yeast GAL4 upstream activating sequence (UAS) (44). To determine whether OsTLP7, and its separate domains, can function as transcriptional activators, we therefore fused GAL4 to the N-terminus of OsTLP7, OsTLP7-FboxD or OsTLP7-TLD. These constructs were cloned within a RUBY reporter backbone in which betalain synthesis is driven under the control of the GAL4-UAS promoter (46). To observe whether Tie1 binding impacts betalain production, GAL4-tagged OsTLP7 constructs were co-expressed in *N. benthamiana* with either AVR-Pii or Tie1. Although GAL4-OsTLP7-FboxD demonstrated transcriptional activation activity, neither full-length GAL4-OsTLP7 nor GAL4-OsTLP7-FboxD demonstrated significant differences in the intensity of betalain production between co-expression with AVR-Pii or Tie1 (Fig. 4A). However, co-expressing GAL4-OsTLP7-TLD with Tie1 significantly increased levels of betalain production compared to co-expression with AVR-Pii (Fig. 4A and S12A). We hypothesise that Tie1 re-localising the OsTLP7-TLD to the nucleus permits the host protein to function as a transcriptional activator. To further investigate the impact of re-localisation on transcriptional activation, the GAL4-OsTLP7-TLD^PIP2^ was co-expressed with AVR-Pii or Tie1 (Fig. 4B and S12B). Compared to OsTLP7-TLD, OsTLP7-TLD^PIP2^ produced significantly higher levels of betalain when co-expressed with Tie1. However, there was no significant difference found between OsTLP7-TLD and OsTLP7-TLD^PIP2^ when co-expressed with AVR-Pii, suggesting that abolishing PIP_2_ binding may not be sufficient to induce transcriptional activation activity. However, Tie1 binding is adequate to induce transcriptional activation of OsTLP7-TLD.

**Figure 4:**
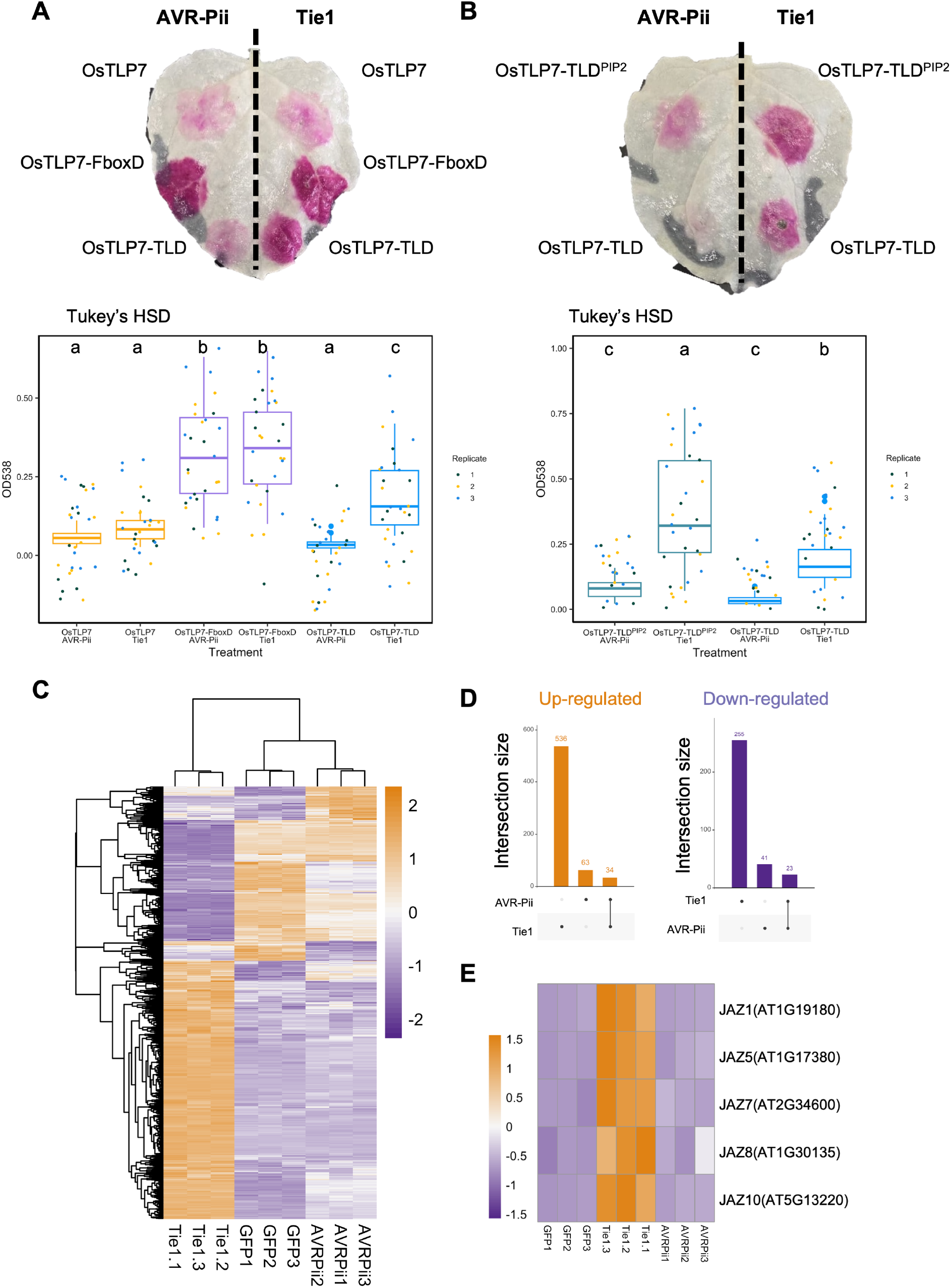
Tie1 induces OsTLP7-TLD transcriptional activation activity and upregulates suppressors of Jasmonic acid signalling. (A) Top, Representative images of *N. benthamiana* leaves 4 dpi after infiltration and after ethanol destaining. Bottom, Quantification of betalain production presented as box plots, with each dot indicating an individual data point and coloured based on replicate number. Letters indicate the statistical significance of each condition using Tukey’s HSD (A, B: p = 2^-16^). The assay shows a significant increase in betalain production when GAL4-OsTLP7-TLD is co-expressed with Tie1-HA compared to meGFP-AVR-Pii-HA. There is no significant difference in betalain production between Tie1-HA and meGFP-AVR-Pii-HA co-expression for GAL4-OsTLP7 and GAL4-F-box domain. (B) As for (A), except the assay shows a significant increase in betalain production when GAL4-OsTLP7-TLD^PIP2^ or GAL4-OsTLP7-TLD is co-expressed with Tie1-HA, compared to GAL4-OsTLP7-TLD with meGFP-AVR-Pii-HA, but not when GAL4-OsTLP7-TLD^PIP2^ is compared to GAL4-OsTLP7-TLD (both with meGFP-AVR-Pii-HA). (C) Hierarchical clustering of significant differentially expressed genes (DEGs) (adjusted *P*-value < 10^−3^, log fold change (|LFC| ⩾ 2) identified in *Arabidopsis thaliana* transgenic lines expressing GFP, GFP-Tie1, or GFP–AVR-Pii under the control of the 35S promoter. Heatmap values represent variance-stabilized, row-centred expression counts. (D) UpSet plots showing shared and specific differentially regulated genes in *A. thaliana*. (E) Heatmap of DEGs showing upregulation of JAZ (Jasmonate ZIM-domain) genes. Numbers below each line indicate replicate number. Expression values are displayed as variance-stabilized, row-centred counts.

Since we observed that Tie1 can bind to TLP proteins from both rice (a monocot) and *N. benthamiana* (a dicot), we investigated the effects of Tie1 overexpression in *A. thaliana*, given the genetic tools available in this model plant. To this end, we generated *A. thaliana* lines stably overexpressing Tie1-GFP, alongside GFP-AVR-Pii and GFP as controls. T₂ generation Tie1-GFP Arabidopsis plants exhibited a pronounced dwarf phenotype accompanied by leaf curling, whereas GFP-AVR-Pii and GFP transgenic lines developed normally (Fig. S13).

To understand the molecular basis of how Tie1 manipulates *A. thaliana* development, we performed RNA-sequencing (RNA-seq) on stable Tie1-GFP lines and compared their transcriptomes with GFP-AVR-Pii and GFP controls. The Tie1-GFP lines displayed a distinct transcriptional profile relative to both controls (Fig. 4C and S14). Differential expression analysis (log2 fold change [|LFC|] ≥ 2 and adjusted *P* < 10^−3^) revealed a substantially higher number of differentially expressed genes (DEGs) in Tie1 (792) than in AVR-Pii (104), with only 57 genes shared between the two (Fig. 4D). Gene Ontology (GO) enrichment analysis of Tie1-regulated genes indicated an overrepresentation of biological processes among upregulated DEGs, including “response to jasmonic acid”, “response to fatty acid”, “response to wounding”, and 17 other GO categories (Fig. S15). Among the upregulated DEGs, five important JA signalling repressors, including JAZ1 (AT1G19180), JAZ5 (AT1G17380), JAZ7 (AT2G34600), JAZ8 (AT1G30135), and JAZ10 (AT5G13220), were highly expressed, suggesting the suppression of JA signalling (Fig. 4E). In total, 23 direct or indirect JA repressor protein genes were upregulated in the Tie1-overexpression background (Fig. S16). GO enrichment analysis also indicated significant downregulation of photosynthesis-related terms which might serve as a pathogen infection strategy (Fig. S15). Further, we observed downregulation of circadian clock-related genes, including LHY (At1g01060), TIC (At3g51240), and At3g51238. Several important auxin-related genes were also downregulated, including SAUR1 (At4g34770), SAUR5 (At4g34810), SAUR15 (At4g38850), SAUR33 (At3g61900), and AXR3 (At1g04250). When considered together, these transcriptomic data suggest that Tie1 may suppress JA signalling, potentially perturbing hormonal homeostasis. Such mis regulation likely contributes to the dwarf and leaf-curling phenotypes observed in Tie1-GFP lines.

## DISCUSSION

To date, SUSS effectors have predominately been studied in an immune receptor recognition context, such as the well characterised MAX effectors that largely interact with HMA proteins (18, 19, 47) and the ZiF family effector AVR-Pii that binds host protein OsExo70F2/F3 (22, 23, 48). However, how sequence diversification of SUSS effectors results in neofunctionalization to promote virulence is less well studied. ZiF effectors can interact with diverse host targets, such as MoHTR1 which directly binds to promoter DNA modulating immune-related gene expression (26) and has recently been proposed to interact with OsSR45 to manipulate alternative splicing of immunity-associated transcripts (25). However, more examples of SUSS effectors interacting with host targets will broaden our understanding of how plant pathogen effectors diversify to promote infection. In this study, we expand the known functional landscape of the ZiF effector family. We identify host Tubby-like F-box proteins as a target of the ZiF effector Tie1 and show how this impacts protein localisation and alters transcriptional activation activity. Collectively, this study supports the concept that SUSS effectors can neo-functionalise to manipulate diverse host targets associated with plant immunity (25, 26).

We initially identified Tie1 as an effector localised to the host delivery system of *M. oryzae* (the BIC) upon infection and demonstrated an interaction with TLPs from *N. benthamiana* and rice. Tubby-like proteins are well conserved (32); and Tie1 likely interacts with TLPs across diverse plant species. Consistent with this, Tie1 homologues from *M. oryzae* strains isolated from diverse host plants all interact with OsTLP7. Targeting a conserved protein family may therefore provide *M. oryzae* with the flexibility to infect multiple hosts and potentially facilitate host transitions (49). Effectors such as Tie1 that target conserved host proteins represent promising candidates for future engineering projects aiming to restrict pathogen spread.

Virulence assays of Tie1 knockout *M. oryzae* Guy11 strains did not show a significant defect on disease progression on rice, most likely due to functional redundancy of *M. oryzae* effectors. Other effectors secreted by Guy11, for example, may also target TLPs, or a conserved downstream pathway may be regulated by diverse TLPs. Although TLP proteins are conserved throughout plant species their overarching roles remain poorly characterised, although they have been linked to immunity in some cases (34, 50). Here, we observed that the localisation of the OsTLP7-TLD changes upon Tie1 binding, which could present a pathogen strategy to uncouple transcriptional regulation of the host defense response to infection. Precedent of strategies based on pathogen effector relocalisation of host targets to promote virulence include Pwl2, which re-localises the host protein HIPP43 away from plasmodesmata (20). Moreover, the Phytophthora RxLR effector Pi03192 targets two NAC transcription factors (51), re-localising these proteins from the endoplasmic reticulum to the nucleus, preventing nuclear accumulation and impeding transcriptional activity. By contrast, our data supports Tie1 in promoting re-localisation of a host transcriptional regulator to the nucleus to promote susceptibility. Additionally, TaTLP5 has recently been shown to promote ubiquitination-dependent degradation of the susceptibility factor Catalase1, that reduces ROS production (52). This demonstrates the broad regulatory role that TLPs have in plants.

Using Alphafold3, we identified a putative interaction interface between Tie1 and OsTLP7-TLD. A single point mutation (OsTLP7-TLD^F336E^) perturbed the localisation of OsTLP7-TLD and its relocalisation to the nucleus on co-expression with Tie1. This suggests Tie1 targets a region of OsTLP7-TLD important for OsTLP7 function. Although the identified putative interaction interface is distal to the PIP_2_ binding site we consistently observed a correlation between the effect on the OsTLP7-TLD of PIP_2_ binding mutants and Tie1 binding. While not direct evidence, based on these observations, we hypothesise that Tie1 interaction disrupts PIP_2_ binding to the OsTLP7-TLD. Tie1 may manipulate the localisation of OsTLP7 by either recruiting a phospholipase C, or other host proteins to promote re-localisation, or the binding of Tie1 induces a conformational change in the OsTLP7-TLD to release PIP_2_. The molecular mechanism of how Tie1 interferes with PIP_2_ binding requires further study.

One remaining question is how does Tie1 binding to TLPs induce the transcriptional differences observed in the *Arabidopsis* stable transgenic lines? We found that the Tubby-like domain of OsTLP7 acts as a transcriptional activator when co-expressed with Tie1. Our data also demonstrates that stable overexpression of Tie1 in *Arabidopsis* induces transcriptional changes, including an upregulation of JAZ genes typically associated with immune suppression. Transcriptional differences observed from the transgenic lines may therefore be a result of Tie1 manipulating AtTLPs, or similar to MoHTR1, Tie1 could directly bind DNA to modulate transcription, although the latter seems less likely (26, 44). Previously, OsTLP2 was shown to regulate expression of the OsWRKY13 gene, which has been implicated in immunity (53). However, the contribution of each TLP protein to transcriptional regulation remains to be defined. Nevertheless, as TLPs are characterised multifunctional regulatory proteins, they are an ideal target for pathogens to manipulate and promote disease.

In summary, we identify plant host Tubby-like proteins (TLPs) as an under-studied protein family targeted by the *M. oryzae* ZiF effector, Tie1. While knowledge of how Tie1 might promote pathogen virulence requires further investigation, our study suggests a role for TLPs in mediating plant immune signalling pathways. Nevertheless, the conservation of the Tie1 homologues across diverse *M. oryzae* strains suggests this effector represents a promising candidate to develop novel resistance strategies against wheat and rice blast disease, for example by immune receptor engineering.

## MATERIALS AND METHODS

### Molecular cloning and transformation of plants

Plasmids were constructed using either In-Fusion cloning (Takara Bio, USA) or the Golden Gate system (54), with modules generated from TSL SynBio. Details of all constructs, including epitope tags, are provided in Supplementary Table 1. DNA modules were obtained via PCR amplification, synthesized as Gblocks (IDT), or acquired from TSL SynBio. Restriction enzymes were from New England Biolabs (Ipswich, MA, USA). Primer sequences and their applications are listed in Supplementary Table 2. Site directed mutagenesis was performed using the Q5^®^ Site-Directed Mutagenesis Kit from New England Biolabs (Ipswich, MA, USA) according to the manufacturers protocol.

*Escherichia coli* bacteria (Stellar) were grown on LB (Lysogeny broth) plates or in liquid media with appropriate antibiotics at 37°C. *Agrobacterium tumefaciens* (GV3101; C58 (rifR) Ti pMP90 (pTiC58DT-DNA) (gentR) Nopaline(pSouptetR)) containing the relevant binary expression vector were grown on LB plates or in liquid media with appropriate antibiotics at 28°C.

*Nicotiana benthamiana* plants were grown in controlled long-day conditions (16-h light (in combination of 2 Philips ter TL-D 58W/840 and Sylvania GRO-LUX F58W/GRO-T8 fluorescent tubes)/ 8-h dark cycle; 22°C) for 4-5 weeks.

Transgenic *Arabidopsis thaliana* plants were generated using the floral dip method(55) with Agrobacteria (strain GV3101) containing the desired expression constructs. Transgenic plants were further selected (kanamycin) and propagated for homozygous seeds Plants were grown under a 16-h photoperiod (12 h light/8 h dark) at 22 °C during the day and 20 °C at night, with a relative humidity of 70%.

### Fungal growth conditions and infection assays

The rice blast fungus *Magnaporthe oryzae* strain Guy11 was maintained on complete medium (CM; 6 g/L yeast extract, 6 g/L acid-hydrolysed casein, 10 g/L sucrose, and 20 g/L agar) at 28 °C under a 12 h light/12 h dark photoperiod. Conidia were harvested from 10-day-old cultures for infection assays.

For pathogenicity assays, conidial suspensions were adjusted to 1 × 10^5^ conidia mL^-1^ and sprayed onto 3-week-old rice plants (*Oryza sativa* cv. CO-39). Inoculated plants were maintained at 28 °C and 80% relative humidity under a 16 h light/8 h dark photoperiod. Disease symptoms were photographed and assessed at 5 days post-inoculation (dpi). Pathogenicity assays were performed in four independent biological experiments with similar results.

To examine effector localisation during infection, a C-terminal GFP fusion of *Tie1f* driven by its native promoter was generated using In-Fusion cloning (Takara Bio, 638909). For co-localisation analyses, constructs expressing the apoplastic effector marker Slp1-mCherry or the cytoplasmic effector marker Pwl2-mCherry were introduced into the *Tie1f*-GFP strain. Rice leaf sheath assays were performed using 3-week-old CO-39 plants. Leaf sheath tissues were inoculated with conidial suspensions (1 × 10^4^ conidia mL^-1^) and incubated at 28 °C. Fluorescence was examined at 26–28 h post-inoculation.

### Generation of *Tie1* deletion mutants

Targeted deletion of *Tie1* homologues was performed using a CRISPR–Cas9-mediated genome editing system. Two single-guide RNAs (sgRNAs) targeting the 5′ and 3′ flanking regions of each gene were designed using CHOPCHOP (https://chopchop.cbu.uib.no/). sgRNAs were synthesized using the EnGen® sgRNA Synthesis Kit (New England Biolabs, E3322), and Cas9 nuclease (EnGen® Spy Cas9 HF1; New England Biolabs, M0667) was used for genome editing.

Donor DNA constructs contained approximately 70-bp homology arms flanking a BAR resistance cassette conferring resistance to glufosinate ammonium. sgRNAs, Cas9 protein, and donor DNA were co-transformed into *M. oryzae* protoplasts by polyethylene glycol (PEG)-mediated transformation. Putative transformants were initially screened by PCR and subsequently confirmed by whole-genome sequencing performed by Novogene. Sequencing reads were mapped to the reference genome and visualized using Integrative Genomics Viewer (IGV) to verify deletion of the targeted loci.

### Bioinformatic online resources used in this study

Signal peptide prediction for the effector proteins from the Tie1 clade effector was carried out using SignalP5.0, (https://services.healthtech.dtu.dk/services/SignalP-5.0/). (56) Alignments and comparative percentage identity analysis were performed using Clustal Omega (http://www.ebi.ac.uk/Tools/msa/clustalo/)(57). The AlphaFold3(37) server was used to generate predictions using three independent seeds. All protein domain boundaries were defined using a combination of InterProScan and AlphaFold3 predictions (37, 58)

### Phylogenetic tree generation

For phylogenetic analysis, we obtained protein sequences from Phytozome database (Oryza sativa v7.0 Phytozome genome ID: 323). We performed protein sequence alignment using MAFFT with auto default parameter. After trimming the alignment with trimAl using gappyout option, we used IQ-TREE (v2.2.2.6) to perform maximum likelihood phylogenetic analysis with bootstrapping (iqtree2 -s Tubby_Rice.fasta -m TEST -alrt 1000 -bb 1000). The ModelFinder option identified ‘Q.insect+F+I+G4” as the optimal substitution model for our dataset. Phylogenetic tree was visualized via TVBOT (https://www.chiplot.online/tvbot.html).

### Transient protein expression in *N. benthamiana* leaves

Transient gene expression in *N. benthamiana* was carried out by co-infiltrating 4-5 week-old plants with *A. tumefaciens* strain GV3101, following established protocols. Each construct was infiltrated at an OD_600_ of 0.3 in Agrobacterium resuspension medium (ARM; 10 mM MgCl₂, 10 mM MES, pH 5.6) supplemented with 150 µM acetosyringone. To suppress post-transcriptional gene silencing, *A. tumefaciens* strain carrying the p19 silencing suppressor from tomato bushy stunt virus was co-infiltrated at an OD_600_ of 0.1. Two leaves per plant were infiltrated, and samples were harvested 48 hours post-infiltration and flash-frozen in liquid nitrogen.

### Pulldown assay from *N. benthamiana* leaves and peptide identification

*N. benthamiana* leaves were transiently transformed as described above with *A. tumefaciens* strain GV3101 carrying the vectors pGG35S::GFP or pGG35S::Tie1-GFP. For this purpose, 2 leaves of 3 individual tobacco plants were infiltrated per construct (6 biological replicates/construct) and after two days whole leave samples were harvested. Following this, all biological replicates per construct were combined to generate single pooled samples and total protein extraction and pulldown assays were carried out. For immunoprecipitation of GFP-tagged proteins 20 µl of GFP-Trap magnetic beads M-270 (ChromoTek) were washed 5 times with extraction buffer GTEN (25 mM Tris pH 7.5, 150 mM NaCl, 1 mM EDTA, 10% v/v glycerol, and 0.1% Tween 20). Proteins were eluted from magnetic beads by boiling at 70 °C for 10 min in 60 µl 2x Laemmli sample buffer. For protein identification 5 µl of the eluted protein fraction were loaded on a SDS gel.

For mass spectrometry analysis, 50 µl of protein sample was mixed with 300 µl of a 1:4 chloroform:methanol solution and incubated on ice for 30 mins. Samples were centrifuged at 15,000 × g for 15 mins at room temperature. The resulting pellet (pellet not visible) was washed with 1 ml methanol, followed by centrifugation at 15,000 × g for 10 mins. This methanol wash step was repeated once more time, followed by a final wash with acetone. After complete drying to remove residual acetone, protein pellets were submitted for mass spectrometry analysis.

### Co-immunoprecipitation and protein immunoblotting

Transient gene expression in *N. benthamiana* was carried out by co-infiltrating 4-week-old plants with *A. tumefaciens* strain GV3101 as described above. Total proteins were extracted from 2 g of tissue using 2 mL GTEN buffer (25 mM Tris-HCl pH 7.5, 150 mM NaCl, 1 mM EDTA, 10%, glycerol, 2%, PVPP, 10 mM DTT, 0.1% Tween-20, and 1x protease inhibitor cocktail (Sigma-Aldrich)). Lysates were clarified by centrifugation at 4,000 × g for 30 mins at 4 °C. For protein pull-downs, 20 µL of M2 anti-FLAG magnetic beads (Sigma-Aldrich) were added to each sample and incubated for 1 hours at 4 °C on a rotary mixer. Samples were washed five times with 1 mL of GTEN buffer (25 mM Tris-HCl pH 7.5, 150 mM NaCl, 1 mM EDTA, 10% [v/v] glycerol, 0.1% Tween-20), and proteins were eluted by heating the beads at 70 °C in 30 µL of 2xSDS sample buffer for 10 mins.

Approximately 15 µL of each eluate was analysed by SDS-PAGE followed by immunoblotting using HRP-conjugated antibodies anti-FLAG (Cohesion Biosciences; 1:3000), anti-GFP (Santa Cruz Biotechnology; 1:3000), and HA-probe (Santa Cruz Biotechnology; 1:5000).

Protein samples were prepared from five leaf discs (8 mm diameter) collected from *N. benthamiana* leaves at 2 days post-agroinfiltration. Discs were homogenized in 100 μL of 2× SDS loading buffer. Proteins were separated by SDS-PAGE and transferred onto polyvinylidene difluoride (PVDF) membranes using the Trans-Blot Turbo Transfer System (Bio-Rad) following the manufacturer’s instructions.

Immunoblotting was performed using the following primary antibodies: HA-probe (Santa Cruz Biotechnology; 1:5000), anti-GFP (Santa Cruz Biotechnology; 1:3000), and anti-FLAG (Cohesion Biosciences; 1:3000). Signal detection was carried out using the Clarity Max Western ECL Substrate (Bio-Rad) and visualized with the ImageQuant LAS 500 imaging system (GE Healthcare). Membranes were stained with Ponceau solution (Sigma-Aldrich, P7170) to confirm equal protein loading.

### Confocal microscopy and image processing

Confocal microscopy images were taken using a LSM 880 from Zeiss with Airyscan and the ZEN (black edition) system 2.3 software. For localisation and co-localisation studies the fluorescence proteins eGFP (488 nm), mRFP (514 nm) and eCFP (458 nm) were excited with an Argon laser, respectively. Emission of eGFP was detected from 489-552 nm, mRFP from 578-632 nm and eCFP from 462-499 nm. Pictures were processed using Fiji, https://imagej.net/software/fiji/downloads(59).

### Transcriptional activation *GAL4 Ruby* assay

To investigate transcriptional activation activity in *N. benthamiana* split GAL4 Ruby assay constructs were used, following the protocol described by Chen *et al*., 2023 (46). Briefly, *A. tumefaciens* strains carrying constructs encoding OsTLP7, the OsTLP7-TLD, OsTLP7-TLD^PIP2^, or F-box domain N-terminally fused to the GAL4 DNA-binding domain (GAL4 BD) under the control of the CaMV 35S promoter were co-infiltrated with constructs expressing the rice blast effector Tie1, or AVR-Pii, each fused to a HA tag under the same promoter. *N. benthamiana* leaves were harvested and imaged after 4 dpi (days post infiltration). To quantify betalain production, harvested leaves were destained with 100% ethanol until all chlorophyll was removed. Once destained a diameter of 0.8 cm was removed from each infiltration site and put into water to extract betalain. Betalain accumulation was then measured using a Grenier F-bottom clear plate in a microplate reader (Clariostar, BMG Labtech) at 538 nm wavelength. All experiments were performed using 10 leaves from 5 different plants and repeated three times.

To confirm protein expression, samples were collected at 3 dpi, flash-frozen in liquid nitrogen, and homogenized. Proteins were extracted by heating the tissue in SDS sample buffer (need recipe) at 70 °C for 10 mins. After centrifugation, supernatants were subjected to SDS-PAGE followed by western blotting. Membranes were probed using anti-Flag-HRP conjugated (Cohesion Bioscience, CSA5714) to detect expression of fusion proteins and anti-HA-HRP conjugated (Invitrogen by Thermo Fisher Scientific, YH379645) for effector proteins.

### RNA isolation from plant material

For mRNA extraction, frozen plant material was homogenized using a mortar and pestle. A total of 2 mg of plant material was used, and mRNA was extracted using the RNeasy Plant Mini Kit for RNA Extraction (Qiagen) according to the manufacturer’s protocol.

### RNAseq (Library preparation and sequencing)

RNAseq analysis was performed by Genewiz (Azenta, Life Science). Briefly, RNAseq library preparation was done by mRNA enrichment, mRNA fragmentation and random priming. Further, first and second strand cDNA synthesis, end repair, 5’ phosphorylation and dA-Tailing, adaptor ligation, PCR enrichment and sequencing. Sequence reads were trimmed to remove possible adapter sequences and nucleotides with poor quality using Trimmomatic v.0.36. The trimmed reads were mapped to the Arabidopsis_thaliana TAIR10.60_ENSEMBL reference genome available on ENSEMBL using HISAT2. Features count function from Subread package was used to obtain raw read counts for downstream analysis. DEGs were identified with DESeq2 following pairwise comparisons between controls. We focused only on DEGs with a strict cutoff (absolute LFC [log2 fold change] ≥ 2 and adjusted P-value ≤ 10−3) for downstream analysis. Heatmaps of differentially expressed genes were generated using the R package pheatmap, and all other plots were produced using custom R scripts. GO term analysis is performed using ShinyGO 0.85.1 with default setting.

## ACKNOWLEDGMENTS

We are grateful to all members of the Banfield Laboratory for valuable discussions. We especially thank Nathan Williams for his careful reading of the manuscript. We further thank Carlo Martins and Gerhard Saalbach (JIC Proteomics), Phil Robinson (Scientific Photographer), Matt Smoker and Jodie Taylor (TSL Tissue Culture and Transformation), and Mark Youles and Liam Egan (TSL SynBio) for their expert assistance and advice throughout this work. We also thank Weibin Ma for assistance with fungal transformation and Vincent Were and Matthew R. Wengler for assistance with mapping whole-genome sequencing reads from the knockout strains.

The authors thank the Biotechnology and Biological Sciences Research Council (UKRI-BBSRC, UK, grants BB/V015508/1, BB/P012574, BBS/E/J/000PR9795, BBS/E/J/000PR9796, BB/Y002997/1 and BB/X010996/1), the BBSRC Norwich Research Park Biosciences Doctoral Training Partnership (grant BB/T008717/1) and the Gatsby Charitable Foundation for funding.

## Author contribution statement

Conceptualisation: I.S., J.W.B., L.H., N.J.T. and M.J.B.; Methodology: I.S., J.W.B., L.H. and K-S. C.; Software: K-S. C.; Validation: I.S., J.W.B., L.H., K-S. C., N.J.T. and M.J.B.; Formal analysis: I.S., J.W.B., L.H. and K-S. C.; Investigation: I.S., J.W.B., L.H. and K-S. C.; Resources: I.S., J.W.B., L.H. and K-S. C., N.J.T. and M.J.B.; Data curation: I.S., J.W.B., L.H. and K-S. C., N.J.T. and M.J.B.; Writing-original draft: J.W.B., I.S., L.H. and M.J.B.; Writing-review and editing: I.S., J.W.B., L.H. and K-S. C., N.J.T. and M.J.B.; Visualisation: I.S., J.W.B., L.H. and K-S. C.; Supervision: N.J.T. and M.J.B.; Project administration: M.J.B.; Funding acquisition: N.J.T. and M.J.B.

## SUPPLEMENTARY INFORMATION

**Figure S1.**
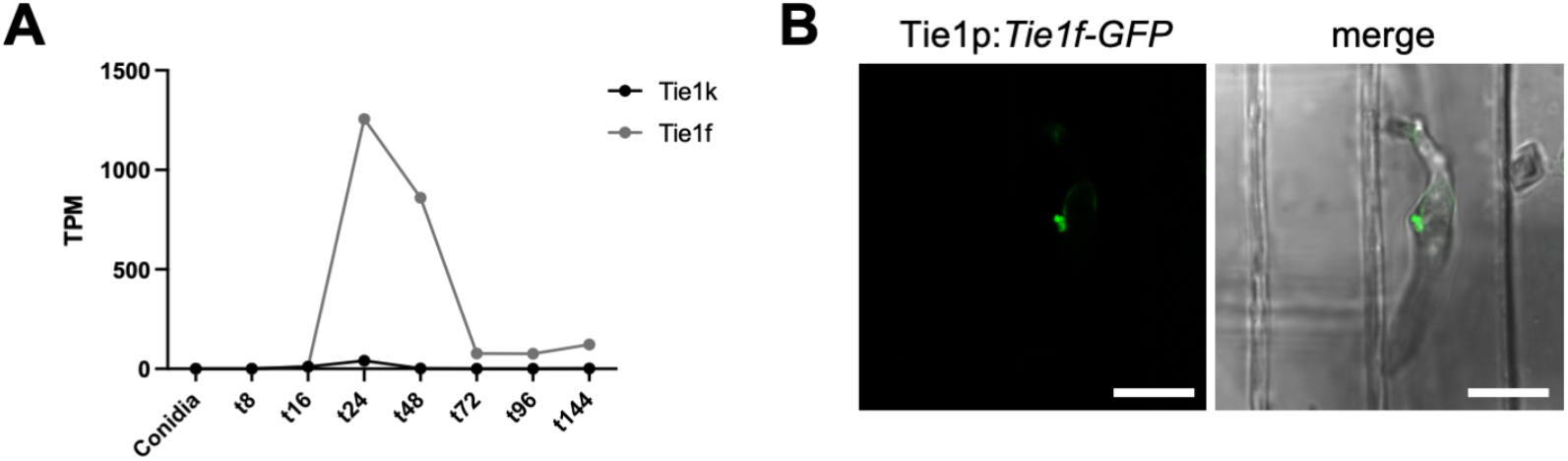
Expression and localisation of Tie1 homologues. (A) Expression profiles of *Tie1k* and *Tie1f* during host infection based on published transcriptome datasets (Yan et al., 2023). Both homologues are expressed during infection, with *Tie1f* showing higher transcript accumulation during the biotrophic stage. (B) Localisation of Tie1f-GFP during rice leaf sheath infection. Tie1f-GFP fluorescence accumulated at the biotrophic interfacial complex (BIC) of invasive hyphae at 26–28 h post-inoculation. Scale bars, 20 μm.

**Figure S2.**
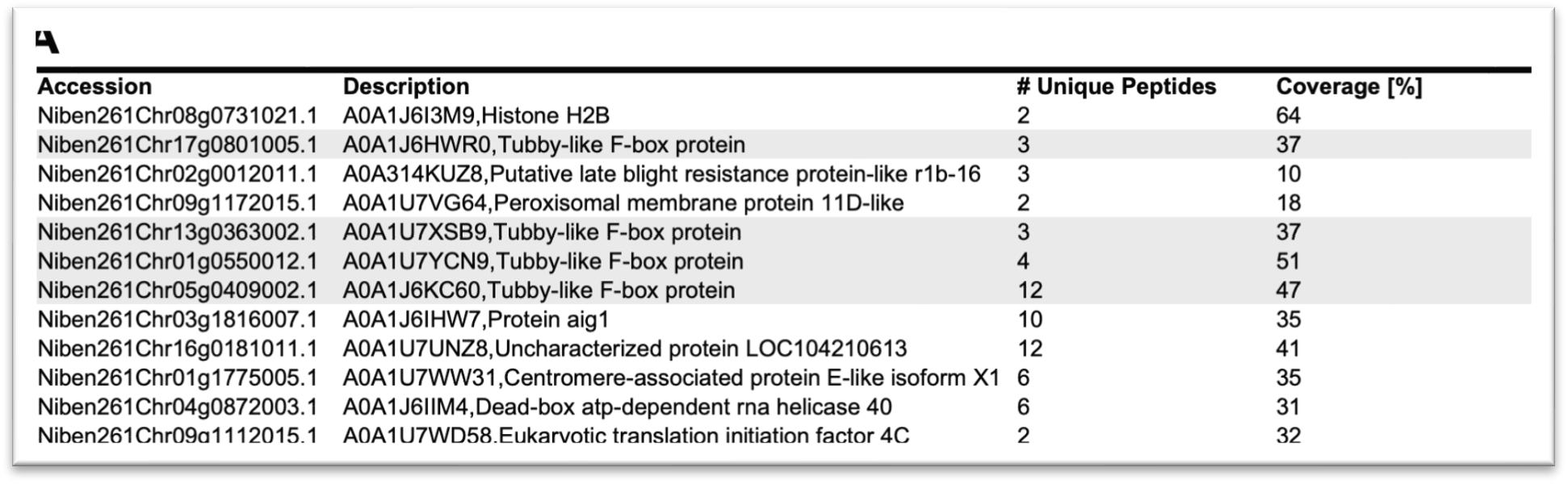
Identification of proteins interacting with Tie1 by IP/MS. Hits from mass spectrometry results following immunoprecipitation in *N. benthamiana*. Column headers include accession numbers (N. benthamiana ID for genes obtained from Sol Genomics Network; N. benthamiana Genome v1.0.1. database), Protein Name, UP (Unique peptide-number of peptides found in the Mass Spectrometry database that are unique for one protein group) and Coverage (percentage of amino acids of the protein sequence covered by identified peptides in the Mass Spectrometry database).

**Figure S3.**
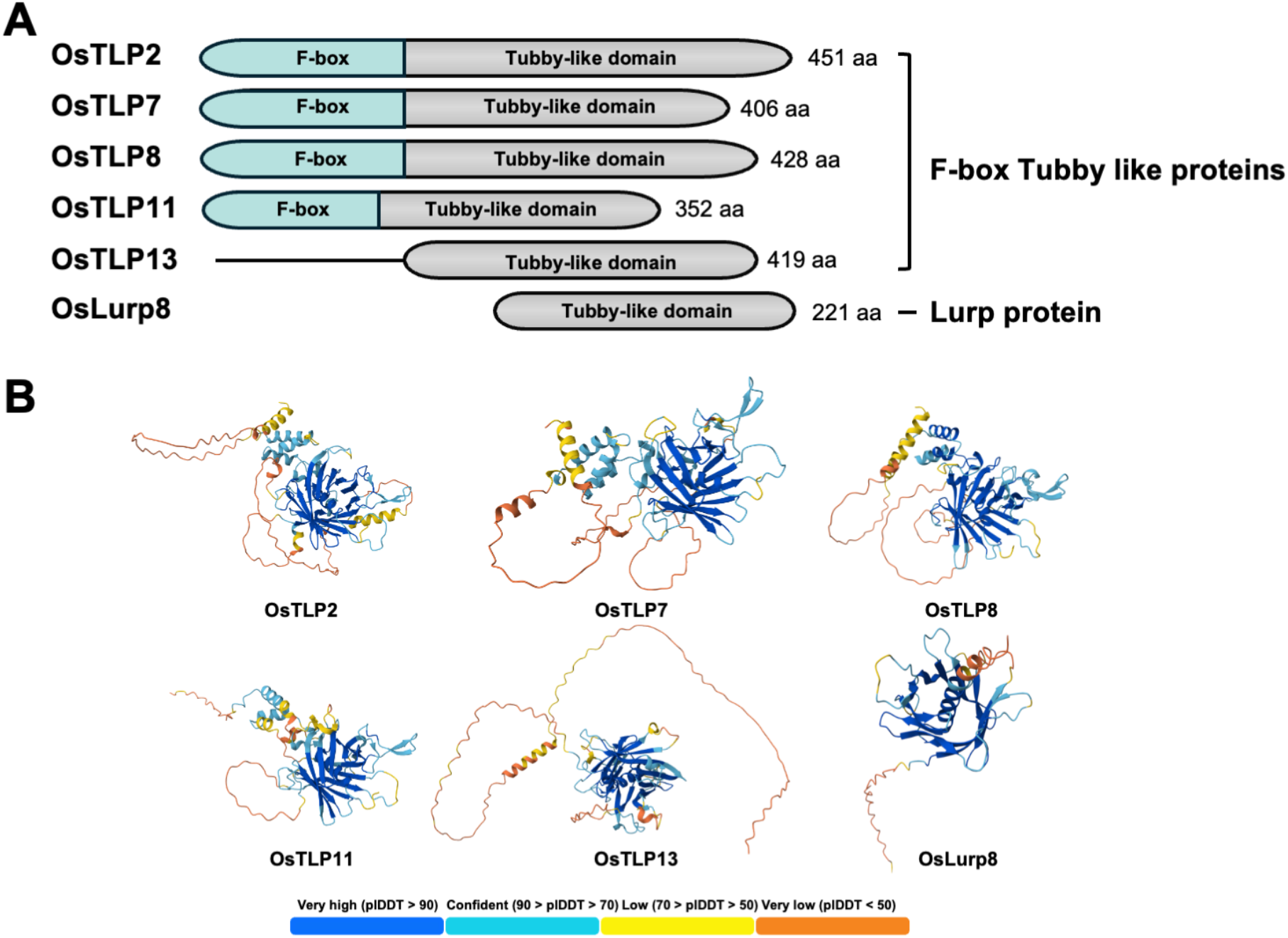
Domain architecture of a subset of *Oryza sativa* Tubby-like domain containing proteins. (A) Protein features of Tubby-like proteins OsTLP2 (LOC_Os08g01290.5), OsTLP7 (LOC_Os07g47110.1), OsTLP8 (LOC_Os02g47640.1), OsTLP11 (LOC_Os05g48670.1), OsTLP13 (LOC_Os02g08310.1), and Lurp8 (LOC_Os11g02750.1). All Tubby-like proteins possess an annotated Tubby-like domain and most have a F-box domain, with the exception of OsTLP13 and Lurp8. (B) AlphaFold3 structural predictions for the full-length proteins OsTLP2 (LOC_Os08g01290.5), OsTLP7 (LOC_Os07g47110.1), OsTLP8 (LOC_Os02g47640.1), OsTLP11 (LOC_Os05g48670.1), OsTLP13 (LOC_Os02g08310.1), and Lurp8 (LOC_Os11g02750.1). AlphaFold3 prediction confidence is represented by a colour gradient from low (orange) to high (blue).

**Figure S4.**
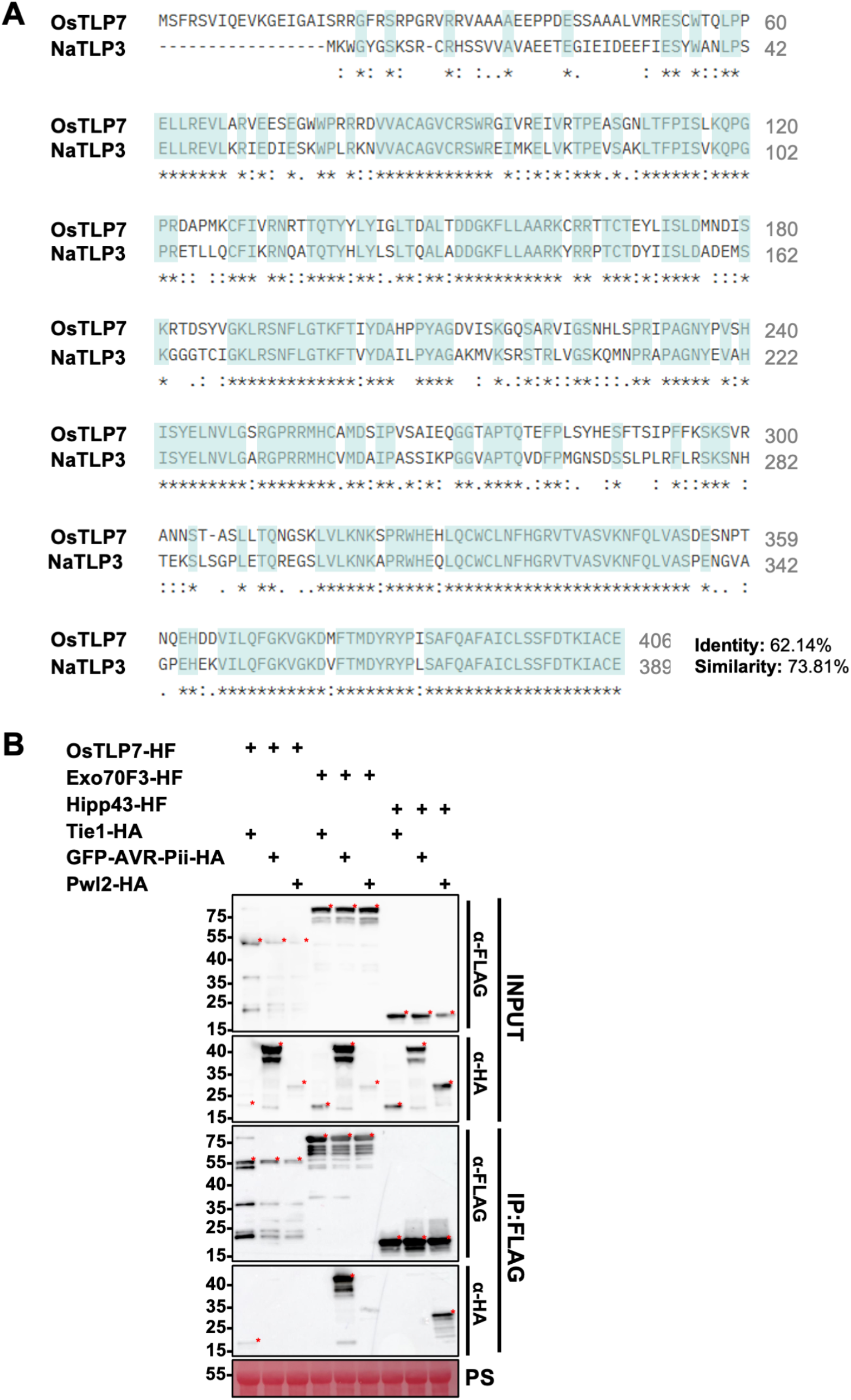
Tie1 specifically binds to OsTLP7. (A) Protein sequence alignment of NaTLP3 with the top hit when queried with NCBI Protein Blast to find similar protein sequences in *Oryza sativa*. Sequence alignment generated using the EMBL-EBI Clustal Omega multiple sequence alignment tool. (B) Co-IP assays were performed in *N. benthamiana* leaves co-expressing OsTLP7-HF, Exo70F3-HF, or Hipp43-HF together with Tie1-HA. These assays reveal a specific interaction between Tie1 and OsTLP7, compared to other *M. oryzae* effector targets Exo70F3-HF (AVR-Pii) or Hipp43-HF (Pwl2). As expected, GFP-AVR-Pii-HA interacted with Exo70F3-HF, but not with OsTLP7-HF or Hipp43-HF, and Pwl2-HA associated with Hipp43-HF, but not with OsTLP7-HF or Exo70F3-HF. Red asterisks indicate the bands of appropriate molecular mass for the expressed protein. Ponceau S staining shows the total amount of protein loaded per lane in the input. The experiment was repeated three times with similar results.

**Figure S5.**
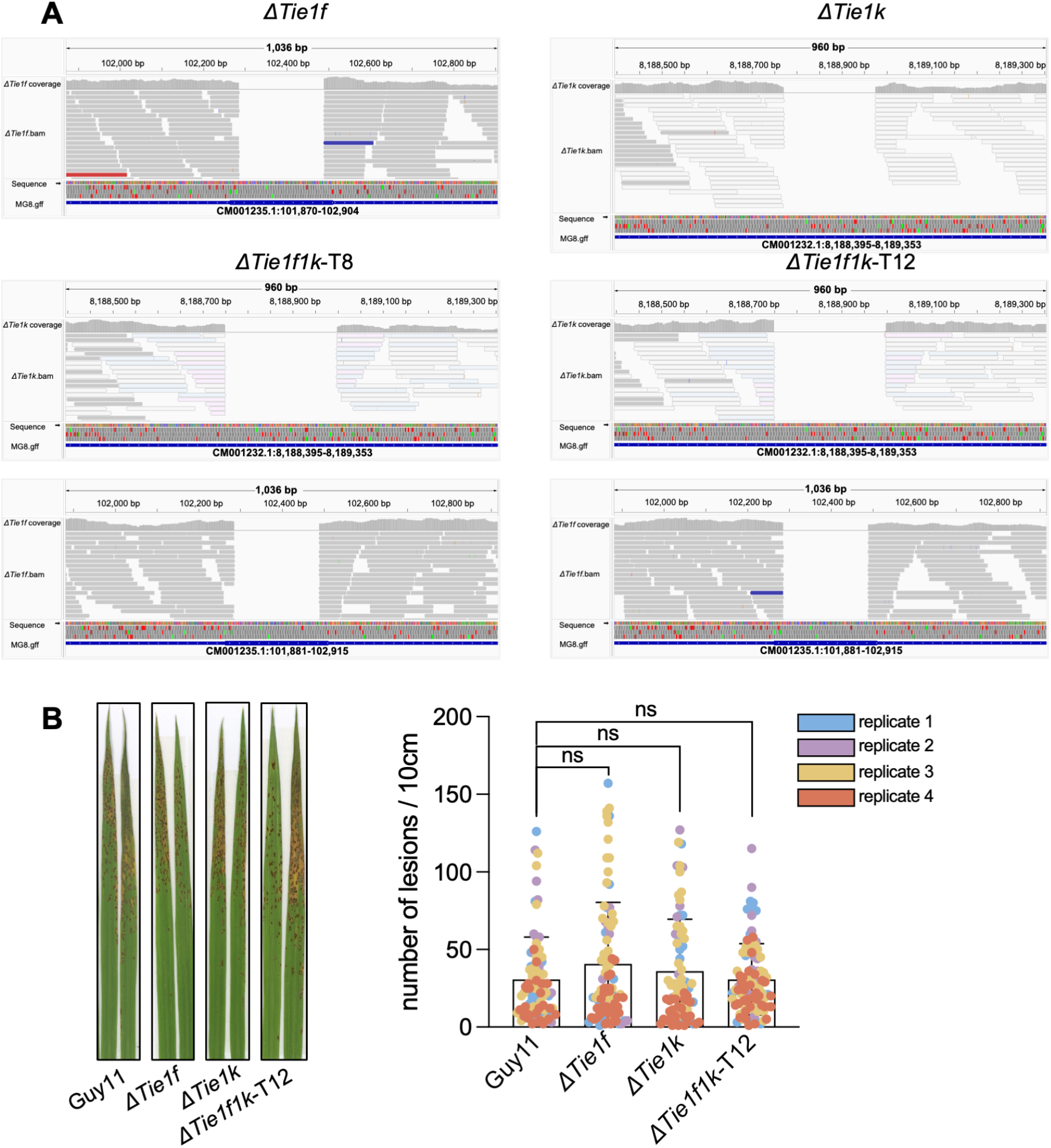
Characterization of Tie1 deletion mutants and pathogenicity assays. (A) Whole-genome sequencing read mapping at the *Tie1k* and *Tie1f* loci. Integrative Genomics Viewer (IGV) screenshots show sequencing read coverage in wild-type Guy11, Δ*tie1k*, Δ*tie1f*, and Δ*tie1k*Δ*tie1f* strains. Loss of read coverage across the targeted regions confirms deletion of the corresponding genes. (B) Pathogenicity assays of wild-type Guy11 and Tie1 deletion mutants on the susceptible rice cultivar CO-39. Rice seedlings were spray-inoculated with 1 × 10^5^ ml^-1^conidial suspensions and photographed at 5 days post-inoculation. No detectable differences in lesion development were observed between wild-type and mutant strains. Representative results from four independent experiments are shown.

**Figure S6.**
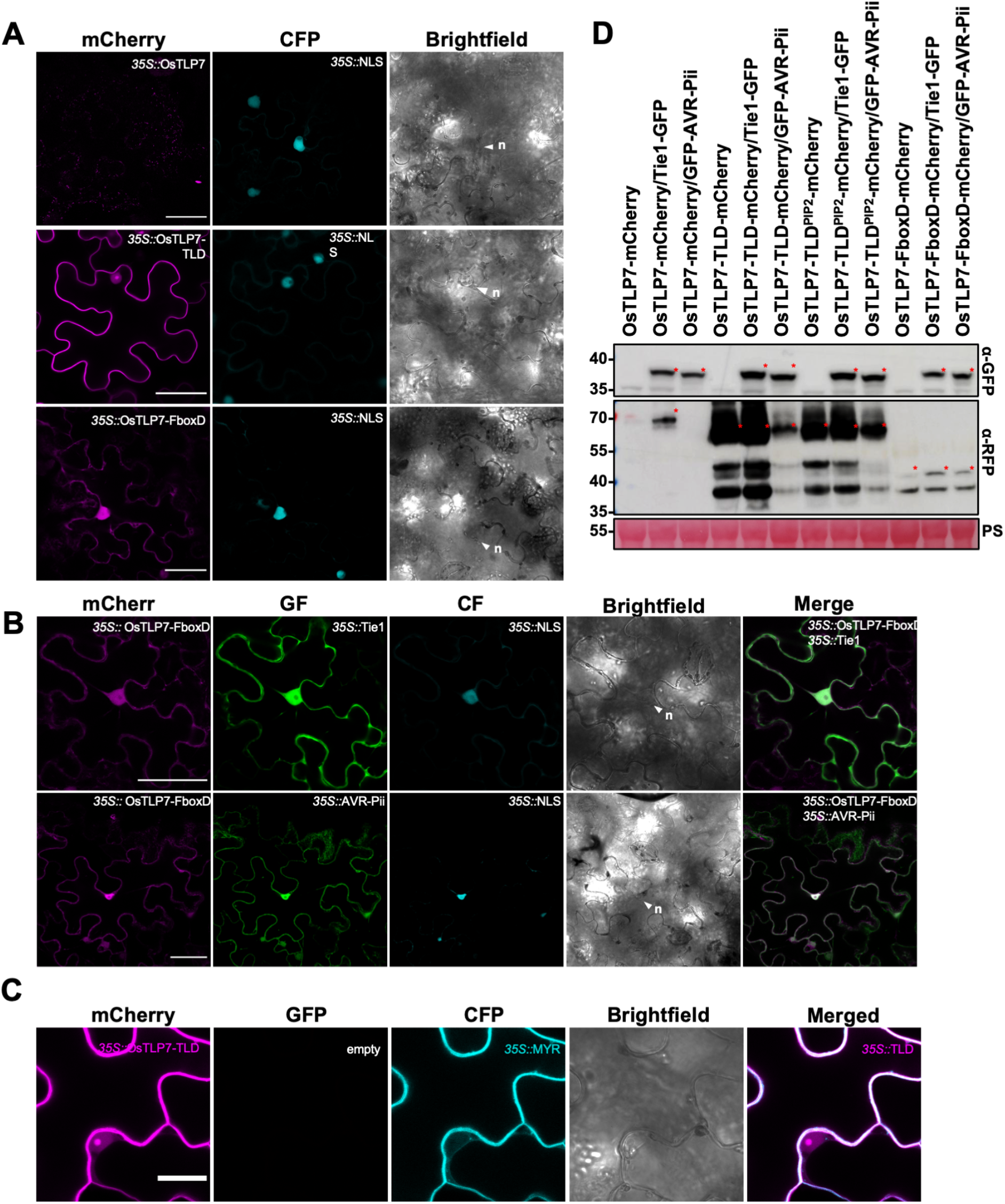
OsTLP7-TLD localises to the nucleolus and plasma membrane while the OsTLP7-FboxD localises to the cytoplasm and nucleus and is not altered by Tie1. (A) Representative images of OsTLP7-mCherry, OsTLP7-TLD-mCherry and OsTLP7-FboxD-mCherry expressed without effectors. CFP-NLS was used to visualize plant nuclei. Nuclei are marked as n; filled white arrowheads indicate nuclei in brightfield images. Scale bar = 50 µm. (B) Representative images of OsTLP7-FboxD-mCherry co-expressed with either Tie1-GFP or GFP-AVR-Pii (control). OsTLP7-FboxD expressed with either Tie1 or AVR-Pii shows a mCherry signal in the cytoplasm and nucleus. CFP-NLS was used to visualize plant nuclei. Nuclei are marked as n; filled white arrowheads in brightfield images. Scale bar = 50 µm. (C) OsTLP7-TLD-mCherry was co-expressed with myristylation(MYR)-CFP, a plasma membrane marker. Clear overlap between OsTLP7-TLD-mCherry and MYR-CFP signals at the plasma membrane is observed. Scale bar = 20 µm. (D) Accumulation of OsTLP7, OsTLP7-TLD, OsTLP7-TLD^PIP2^, and OsTLP7-FboxD fused to mCherry under the control of the 35S promoter in leaf tissue was assessed by western blot using an mCherry antibody. Accumulation of effectors expressed under the 35S promoter was assessed using a GFP antibody. Red asterisks indicate the bands of appropriate molecular mass for the expressed protein. Total protein extracts were visualized by Ponceau (PS) staining to confirm equal loading.

**Figure S7.**
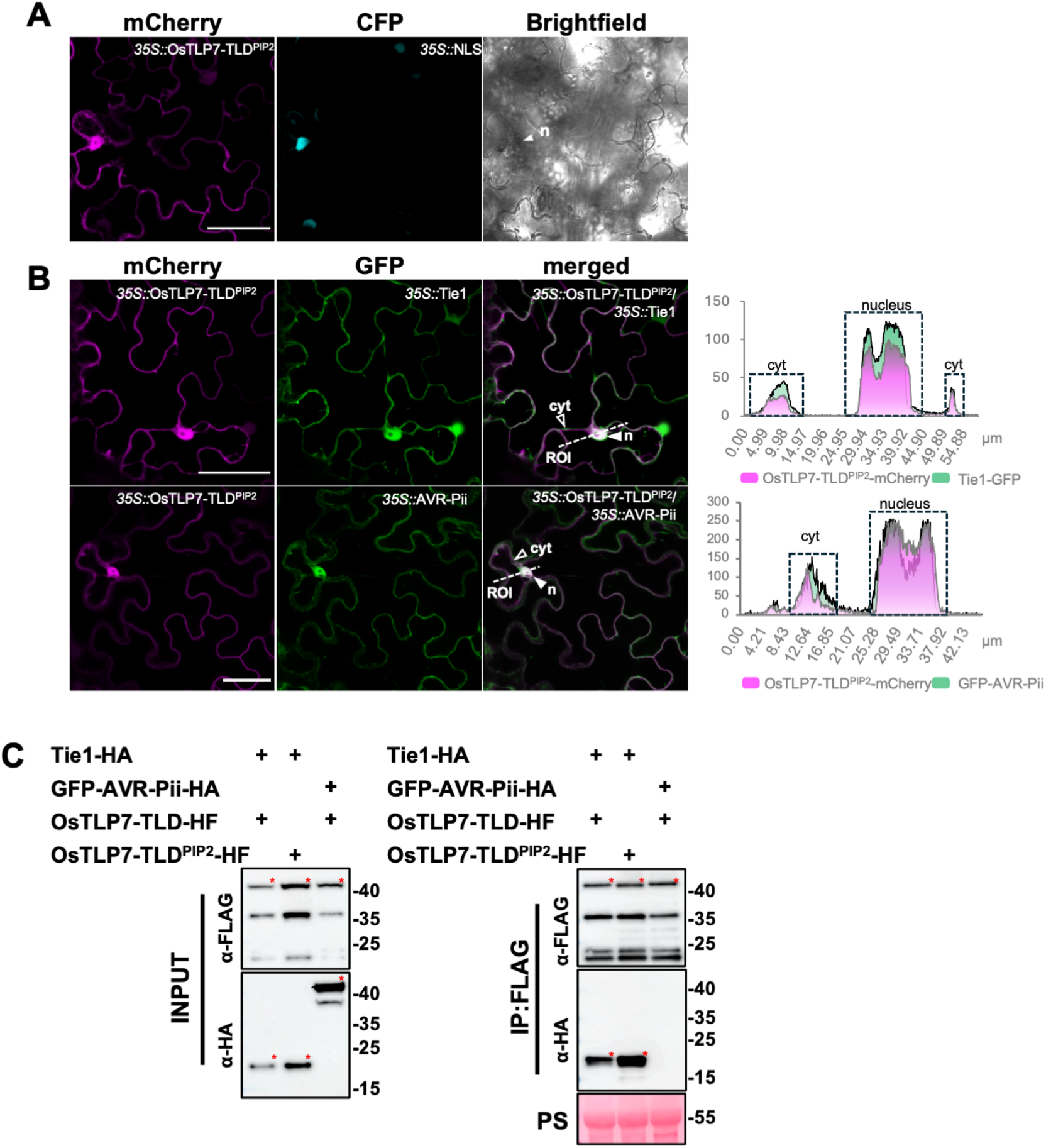
**OsTLP7-TLD^PIP2^ is localised to the cytoplasm and nucleus**. (A) Representative localisation patterns of OsTLP7-TLD^PIP2^. CFP-NLS was used to visualize plant nuclei. Scale bars = 5 µm. (B) Representative localisation patterns of OsTLP7-TLD^PIP2^ co-expressed with Tie1 or AVR-Pii. CFP-NLS was used to visualize plant nuclei. Scale bars = 5 µm.(C) Co-IP assays were performed using OsTLP7-TLD-HF or OsTLP7-TLD^PIP2^-HF co-expressed with Tie1-HA or GFP-AVR-Pii-HA. Tie1-HA specifically interacted with both OsTLP7-TLD and OsTLP7-TLD^PIP2^, whereas no interaction was observed between OsTLP7-TLD and GFP-AVR-Pii. Red asterisks indicate the bands of appropriate molecular mass for the expressed protein. Ponceau S staining confirmed equal protein loading in the input samples. The experiment was repeated three times with consistent results.

**Figure S8.**
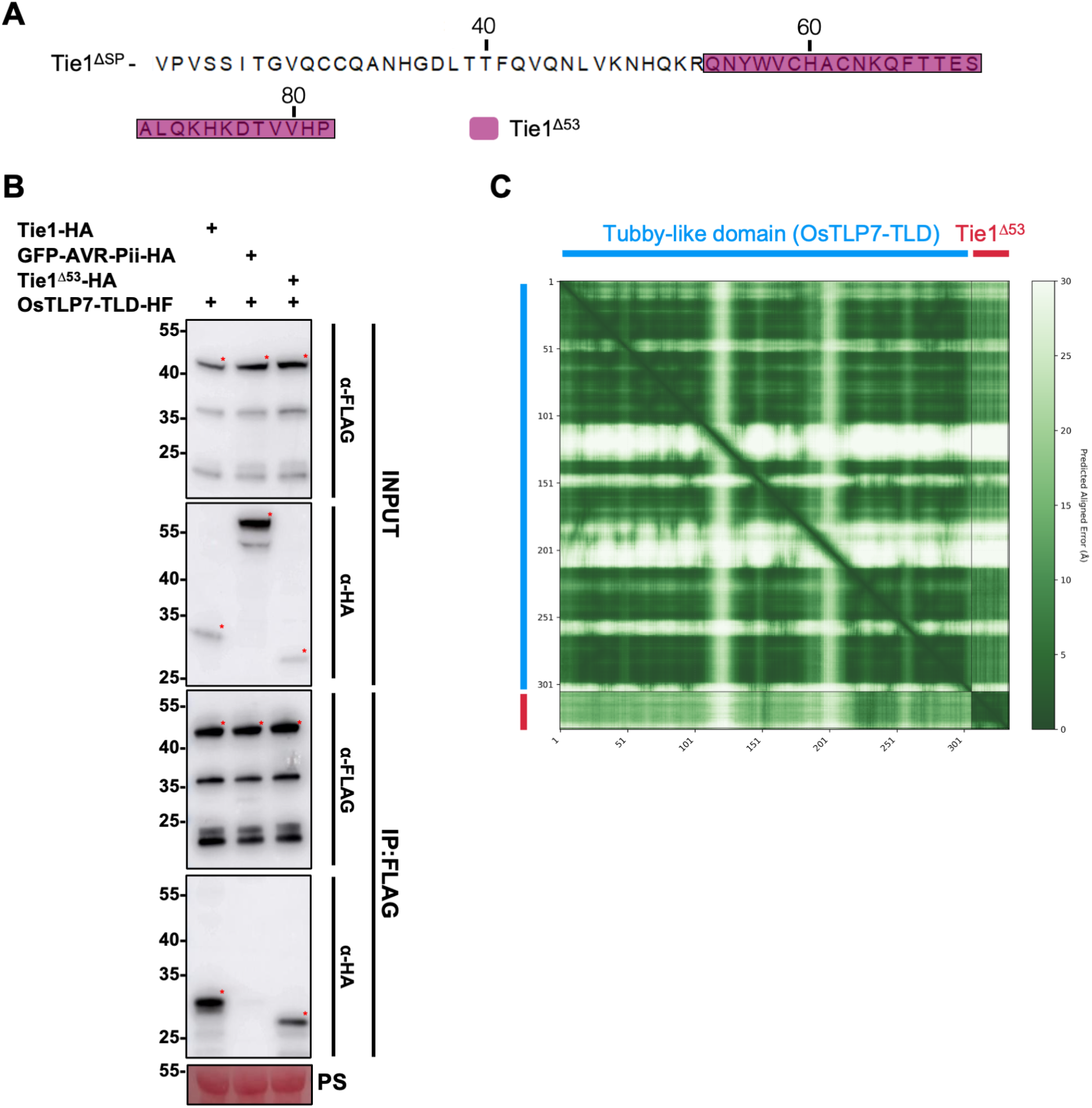
A truncated Tie1 construct retains interaction with OsTLP7-TLD. (A) The sequence of Tie1 without signal peptide, with highlighted regions indicating the truncated Tie1^Δ53^ used in further analysis. (B) Co-IP assays were performed using OsTLP7-TLD-HF co-expressed with Tie1-HA, GFP-AVR-Pii-HA or Tie1^Δ53^-HA. OsTLP7-TLD-HF specifically interacted with both Tie1 and Tie1^Δ53^, whereas no interaction was observed between OsTLP7-TLD and GFP-AVR-Pii. Red asterisks indicate the bands of appropriate molecular mass for the expressed protein. Ponceau S staining confirmed equal protein loading in the input samples. The experiment was repeated three times with consistent results. (C) PAE plot for the interaction of the OsTLP7-TLD with Tie1^Δ53^ for the most confident model presented in all figures.

**Figure S9.**
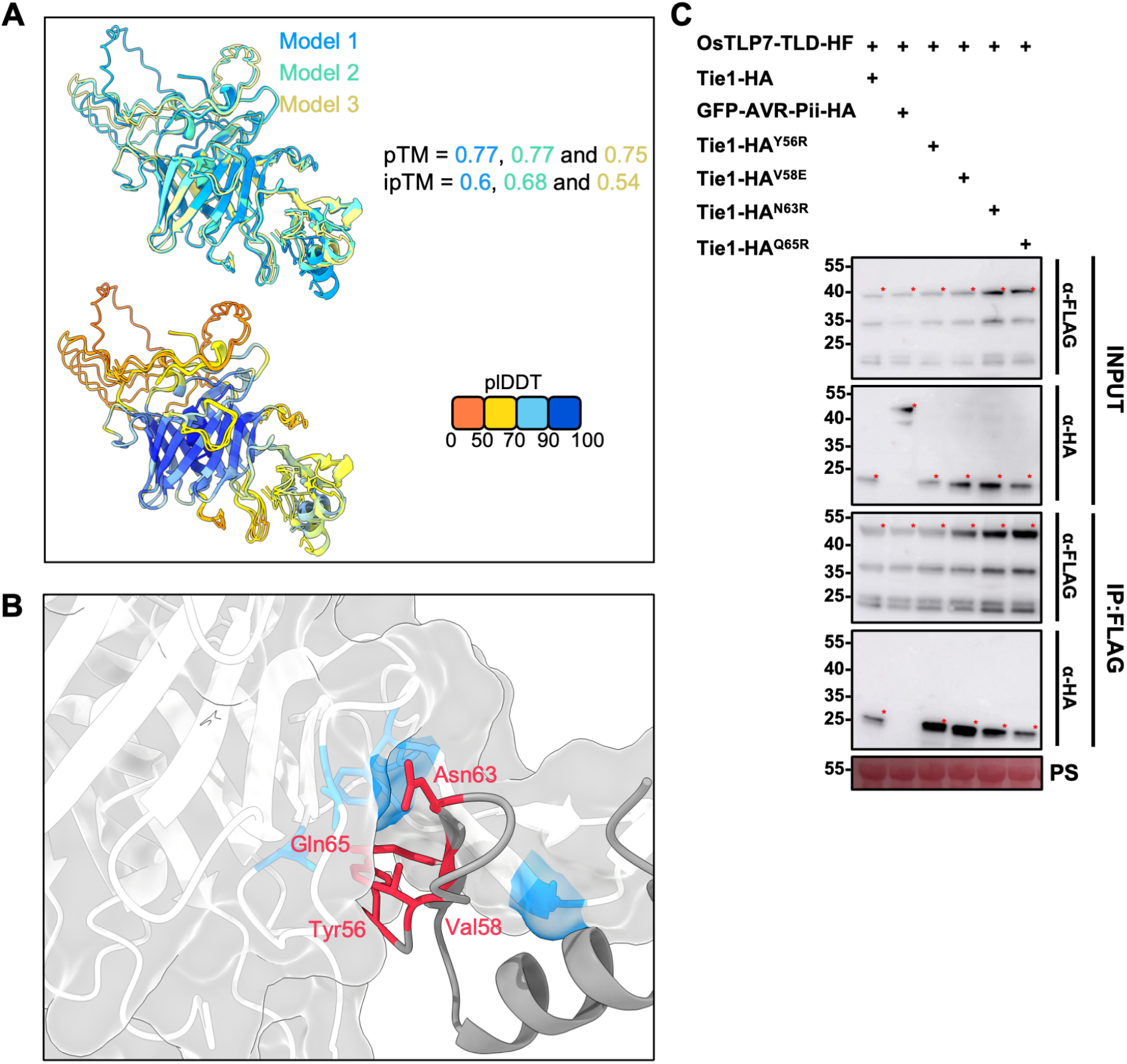
Point mutations of Tie1 based on AlphaFold3 predictions are unable to disrupt interaction with OsTLP7. (A) Repeated AlphaFold3 predictions of OsTLP7-TLD interacting with Tie1^153^ overlayed, with each prediction is coloured according with the pTM and ipTM indicated. The plDDT score for each of the models is shown. (B) A close-up of the interaction interface from the most confident model with residues of Tie1 predicted to be involved in the interaction highlighted. (C) Co-IP assays were performed using OsTLP7-TLD-HF co-expressed with Tie1-HA, Tie1^Y56R^-HA, Tie1^V58E^-HA, Tie1^N63R^-HA, Tie1^Q65R^-HA, GFP-AVR-Pii-HA. OsTLP7-TLD-HF interacted with all Tie1 constructs, whereas no interaction was observed between OsTLP7-TLD and GFP-AVR-Pii. Red asterisks indicate the bands of appropriate molecular mass for the expressed protein. Ponceau S staining confirmed equal protein loading in the input samples. The experiment was repeated three times with consistent results.

**Figure S10.**
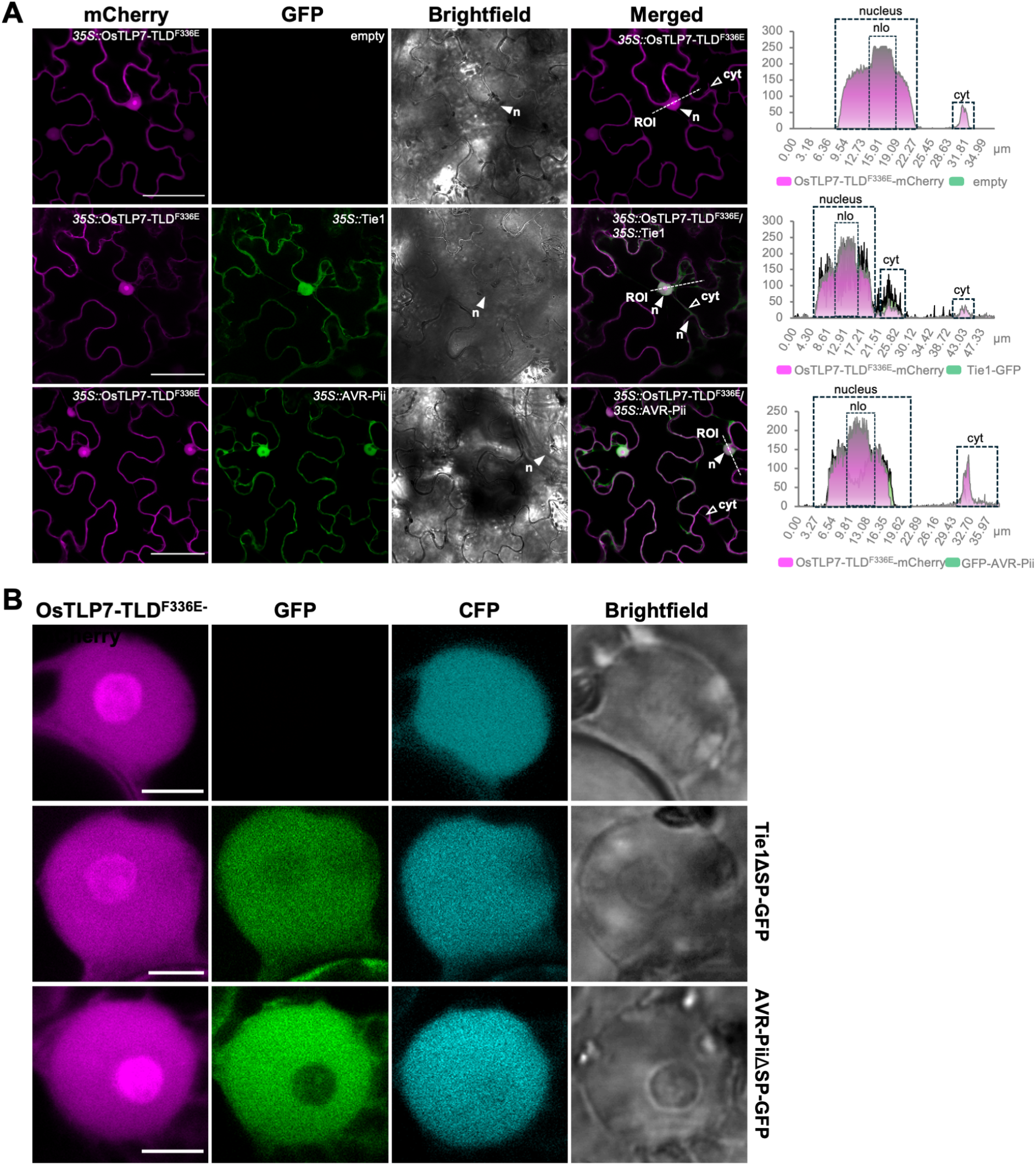
OsTLP7-TLD^F365E^ localisation is disrupted compared to the wildtype but not latered upon coexpression with Tie1. (A) Representative images of OsTLP7-TLD^F336E^-mCherry expressed alone or co-expressed with either Tie1-GFP or GFP-AVR-Pii (control). In all conditions OsTLP7-TLD^F336E^ shows localisation to the cytoplasm and nucleolus. CFP-NLS was used to visualize plant nuclei. Nuclei are marked as n; filled white arrowheads indicate nuclei in brightfield images. Region of interest (ROI, merged images) indicate starting points of transection lines used for fluorescence intensity profiling. Right panels: Fluorescence intensity plots along transection lines for GFP and mCherry channels. Nuclear areas and peripheral cytoplasmic signals are boxed. Maximum fluorescence intensity = 255. Scale bar = 50 µm. (B) Representative localisation patterns of OsTLP7-TLD and OsTLP7-TLD^F336E^ in individual plant nuclei, expressed alone or co-expressed with GFP-Tie1 or GFP-AVR-Pii. CFP-NLS was used to visualize plant nuclei. This panel provides an extended view for Supplementary figure 10A, including the CFP signal and brightfield images. Scale bars = 5 µm.

**Figure S11.**
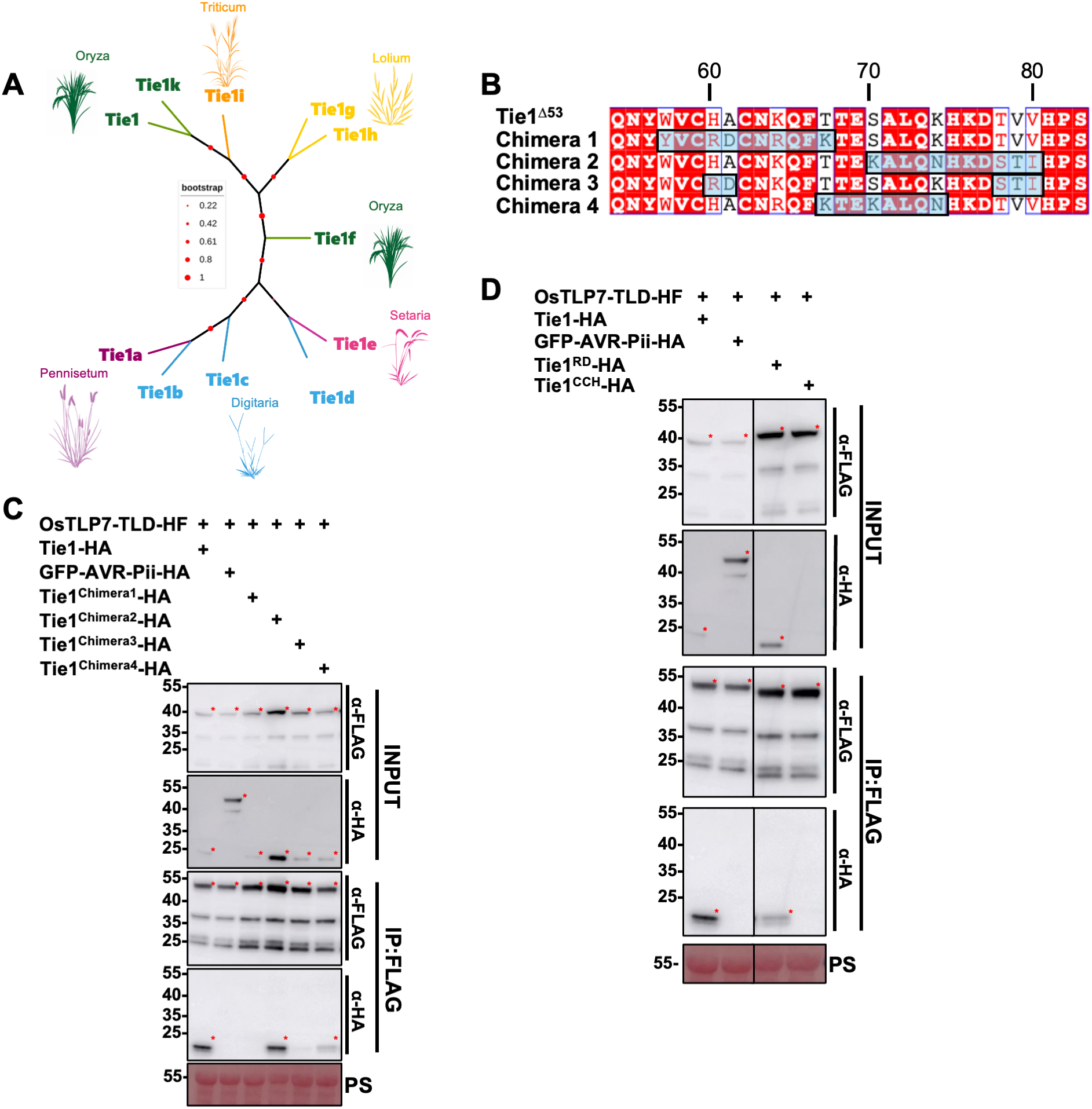
Chimeras of Tie1 and Tie1a reveal a likely interaction interface. (A) Unrooted phylogenetic tree of Tie1 tribe members, with the host species of each Magnaporthe strain indicated (images of plants were generated using ChatGPT4). (B) Sequence alignment of Tie1^Δ53^ against each chimera, with Tie1a with residues highlighted to indicate the regions swapped from Tie1a to Tie1. (C) Co-IP assays of OsTLP7-TLD-HF with four chimeras of Tie1 and Tie1a (all HA-tagged), demonstrating loss of binding in two chimeras (1 and 3). No interactions were detected between OsTLP7-TLD-HF and GFP-AVR-Pii-HA. Red asterisks indicate the bands of appropriate molecular mass for the expressed protein. Ponceau S staining was used to confirm equal protein loading in the input samples. The experiment was repeated three times with consistent results. Western blot images are cropped to exclude unrelated samples in the centre of the image. (D) Co-IP assays were performed using OsTLP7-TLD-HF co-expressed with Tie1-HA, Tie1^RD^-HA, Tie1^CCH^-HA, and GFP-AVR-Pii-HA. OsTLP7-TLD-HF interacted with Tie1 but also with Tie1^RD^. No interaction was observed between OsTLP7-TLD and GFP-AVR-Pii or Tie1^CCH^, however the latter is also missing in the input, suggesting a lack of protein stability. Red asterisks indicate the bands of appropriate molecular mass for the expressed protein. Ponceau S staining confirmed equal protein loading in the input samples. The experiment was repeated three times with consistent results.

**Figure S12.**
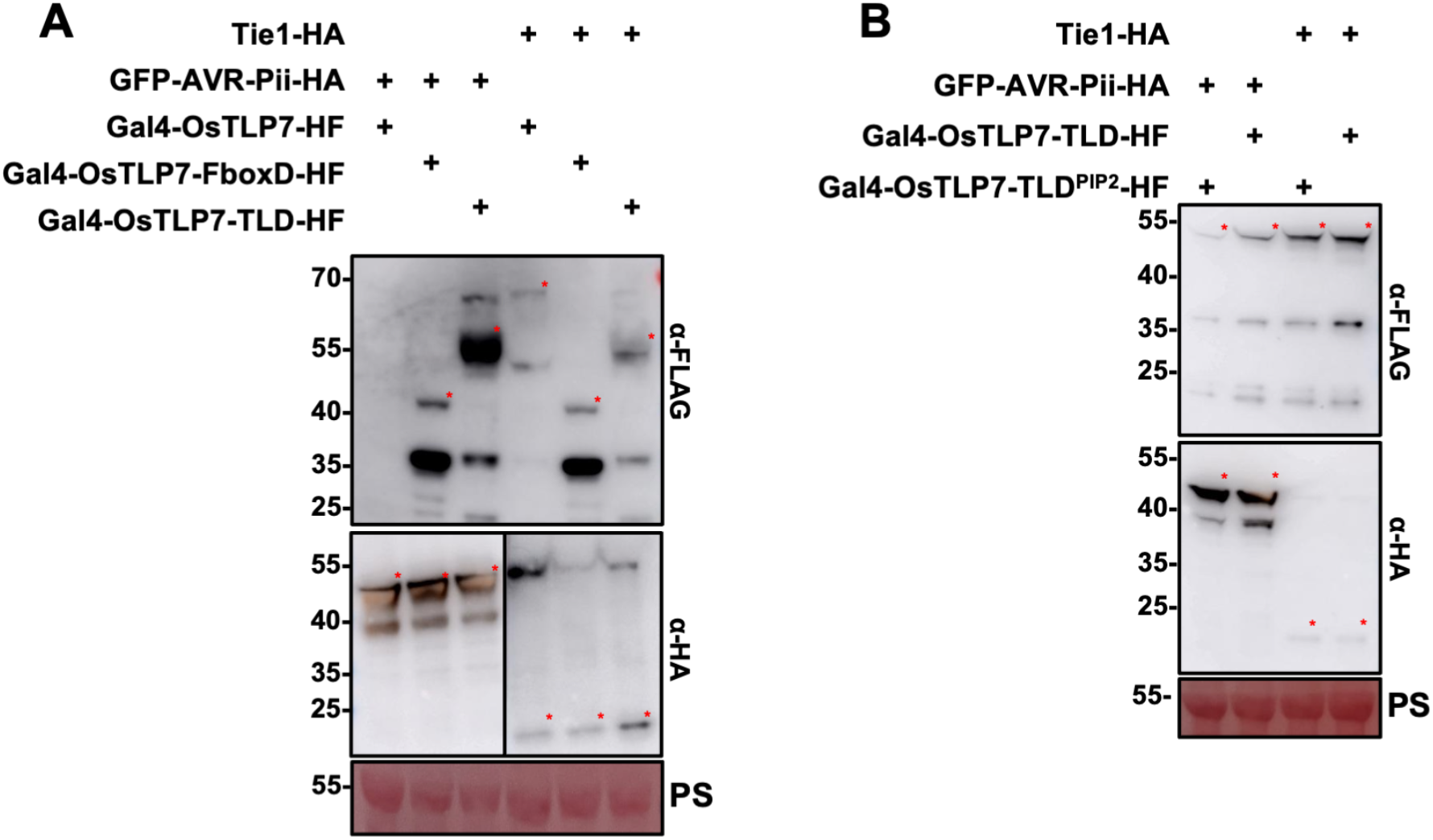
Western blot analysis to confirm protein accumulation of GAL4-tagged constructs. (A) Western blots performed using GAL4-OsTLP7-HF, GAL4-OsTLP7-TLD-HF, or GAL4-OsTLP7-FboxD-HF co-expressed with either Tie1-HA or meGFP-AVR-Pii-HA showing accumulation of all proteins except OsTLP7 when co-expressed with AVR-Pii from the RUBY assays presented in Figure 4. Red asterisks indicate the bands of appropriate molecular mass for the expressed protein. Ponceau S staining confirmed equal protein loading in the input samples. The experiment was repeated three times with consistent results. Western blot images are cropped to prevent overexposure and allow visualisation of all constructs (B) Western blots performed using GAL4-OsTLP7-TLD-HF, or GAL4-OsTLP7-TLD^PIP2^-HF co-expressed with either Tie1-HA or meGFP-AVR-Pii-HA. Showing accumulation of all proteins from the ruby assays in Figure 4. Red asterisks indicate the bands of appropriate molecular mass for the expressed protein. Ponceau S staining confirmed equal protein loading in the input samples. The experiment was repeated three times with consistent results.

**Figure S13.**
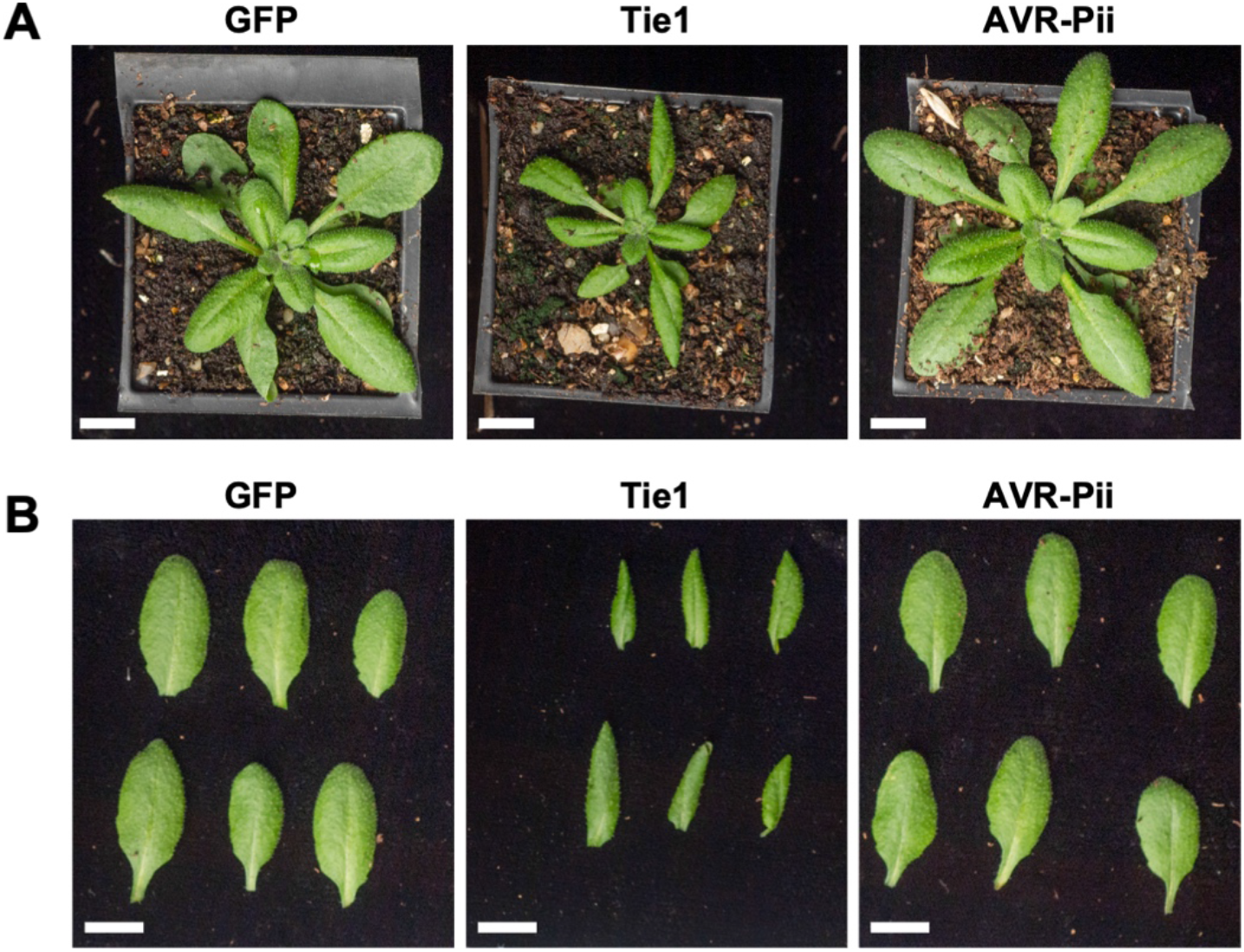
*A. thaliana* stably overexpressing Tie1 exhibit developmental defects. (A) Phenotype of stable *A. thaliana* transgenic lines expressing GFP, Tie1-GFP, and GFP-AVR-Pii. Plants expressing Tie1-GFP display a dwarfed stature and curled leaf phenotype compared to GFP and GFP-AVR-Pii lines. This growth phenotype was consistently observed across three independent transgenic lines. Scale bars = 1 cm. (B) Leaf morphology of stable Arabidopsis thaliana transgenic lines expressing GFP, GFP-Tie1, and GFP-AVR-Pii. Leaves from GFP-Tie1 expressing plants exhibit a curled phenotype compared to GFP or GFP-AVR-Pii lines. This phenotype was consistently observed across three independent transgenic lines. Scale bars = 1 cm.

**Figure S14.**
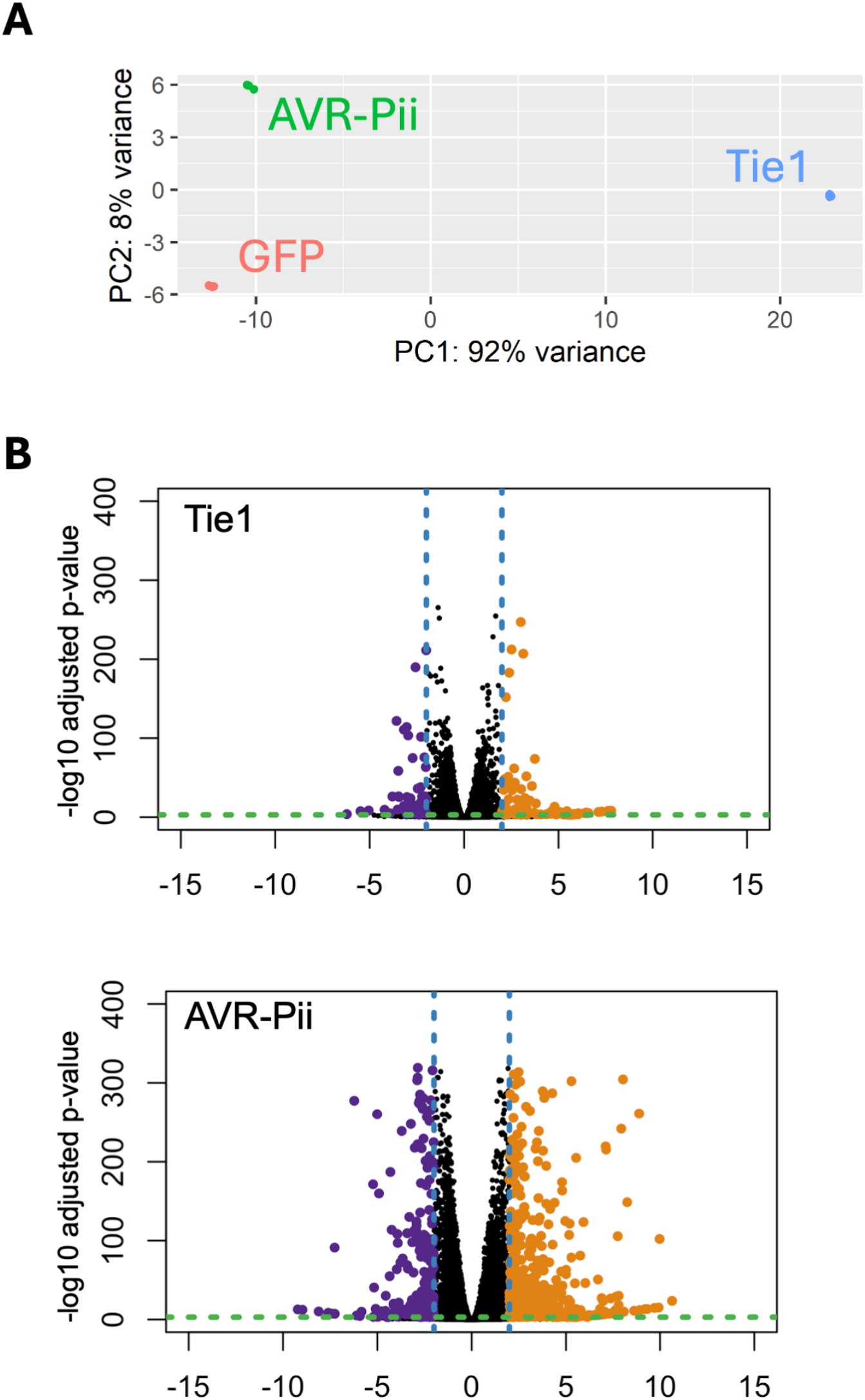
PCA demonstrates transcriptional differences between transgenic *A. thaliana* lines. (A) Principal component analysis (PCA) of *A. thaliana* transcriptome replicates. (B) Volcano plots showing pairwise differential gene expression in stable *A. thaliana* lines expressing GFP-Tie1 and GFP-AVR-Pii compared to GFP control. Genes significantly downregulated are highlighted in purple, significantly upregulated genes are highlighted in green and blue cutoff line at LFC equal to 2.

**Figure S15.**
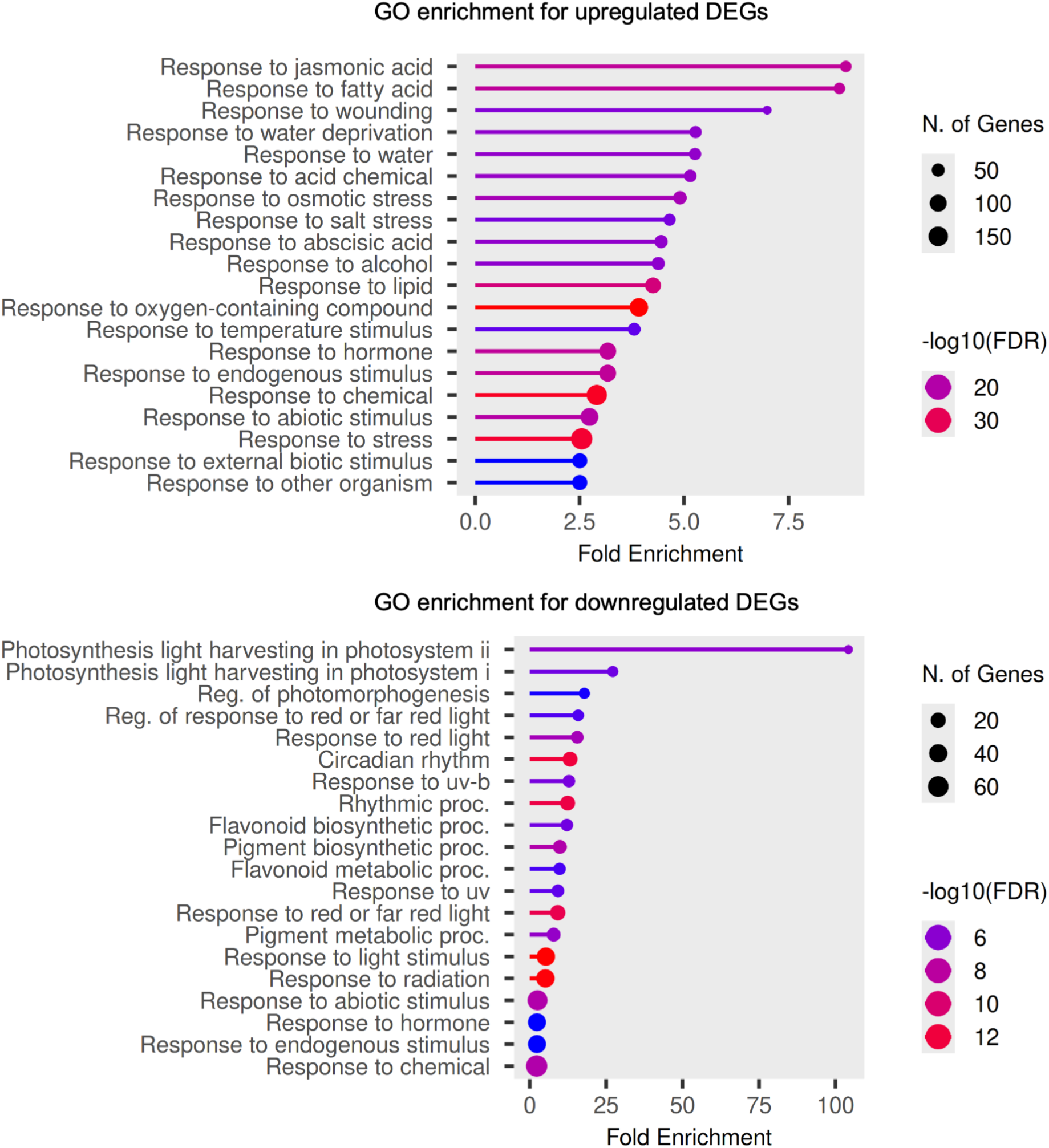
Gene Ontology (GO) term enrichment analysis of DEGs. Comparison of *Arabidopsis thaliana* lines overexpressing GFP-Tie1 versus the GFP control. DEGs identified from the expression analysis were subjected to GO term enrichment using ShinyGO with default parameters. The top 20 enriched biological process terms are displayed, categorized by upregulated and downregulated genes.

**Figure S16.**
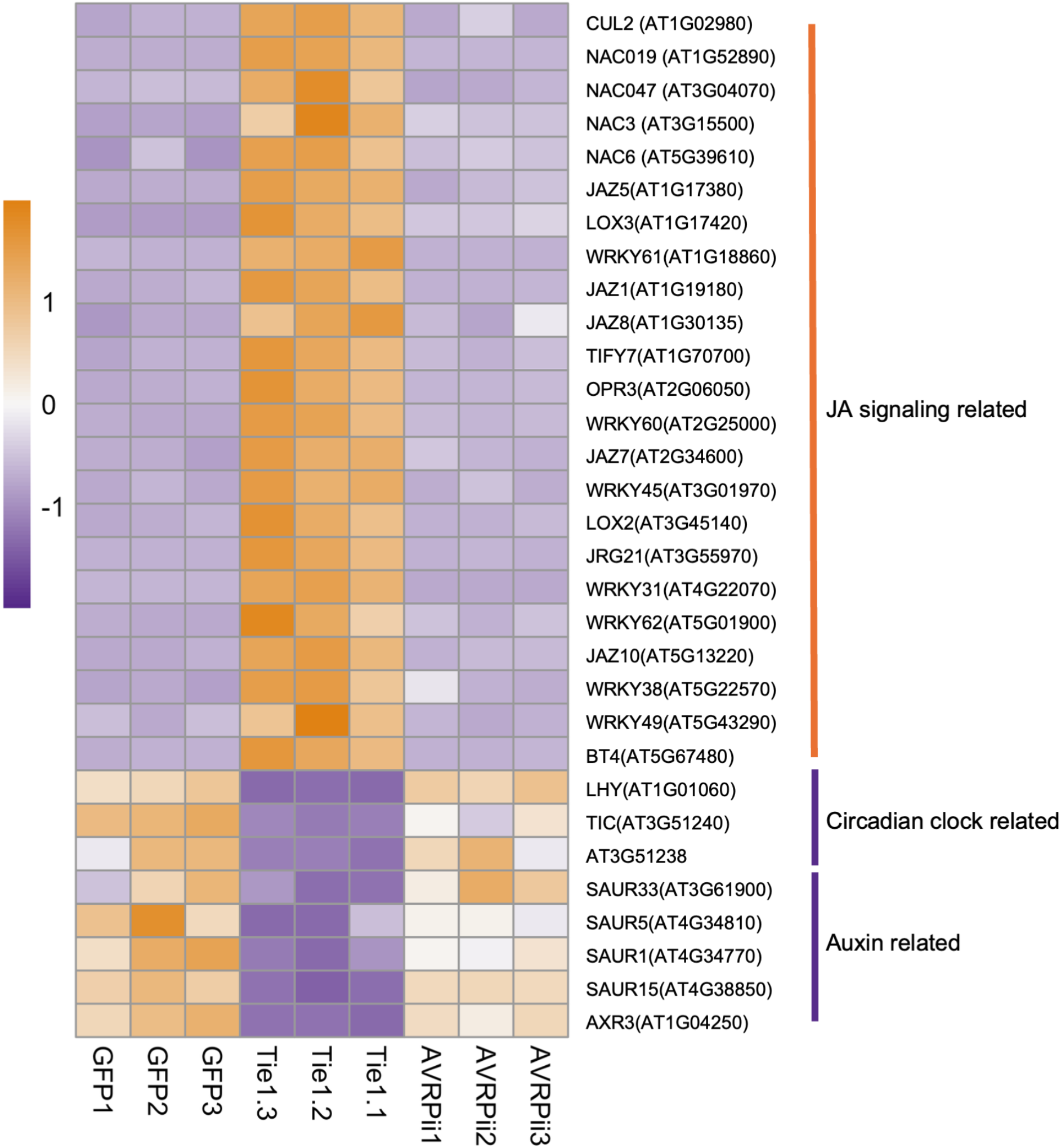
Expression profiles of selected DEGs across targeted functional pathways. Heatmap displaying the log2 fold-change values derived from transcriptome analysis of *A. thaliana* lines overexpressing GFP-Tie1, GFP-AvrPii and GFP controls. Genes were manually curated into three distinct functional categories: the jasmonic acid (JA) signalling pathway, circadian clock-related processes, and auxin-regulated networks.

**Table S1.** Plasmids used in this study.

| **Construct Name** | **RE** | **Plasmid components** | **Resistance** |
| --- | --- | --- | --- |
| GFP-AVR-Pii | Bsal | pICH47742, pICSL13004,<br>pICSL30006, pICH77901 | Carb |
| Tie1-GFP | Bsal | pICH47742, pICSL51277,<br>pICSL50008, pICH41414 | Carb |
| GFP-HA | Bsal | pICH47742, pICSL51277,<br>pISCL30006, pICSL50009,<br>pICH41414 | Carb |
| Tie1-HA | Bsal | pICH47742, pICSL51277,<br>pICSL50009, pICH41414 | Carb |
| GFP-AVR-Pii-HA | Bsal | pICH47742, pICSL51277,<br>pICSL50009, pICH41414 | Carb |
| PWL2-HA | Bsal | pICH47742, pICSL51277,<br>pICSL50009, pICH41414 | Carb |
| OsTLP7-HF | Bsal | pICH47742, pICSL51277,<br>pICSL50001, pICH41414 | Carb |
| Exo70F3-HF | Bsal | pICH47742, pICSL51277,<br>pICSL50001, pICH41414 | Carb |
| Hipp43-HF | Bsal | pICH47742, pICSL51277,<br>pICSL50001, pICH41414 | Carb |
| OsTLP2-HF | Bsal | pICH47742, pICSL51277,<br>pICSL50001, pICH41414 | Carb |
| OsTLP8-HF | Bsal | pICH47742, pICSL51277,<br>pICSL50001, pICH41414 | Carb |
| OsTLP11-HF | Bsal | pICH47742, pICSL51277,<br>pICSL50001, pICH41414 | Carb |
| OsTLP13-HF | Bsal | pICH47742, pICSL51277,<br>pICSL50001, pICH41414 | Carb |
| LURP8-HF | Bsal | pICH47742, pICSL51277,<br>pICSL50001, pICH41414 | Carb |
| OsTLP7-TLD-HF | Bsal | pICH47742, pICSL51277,<br>pICSL50001, pICH41414 | Carb |
| OsTLP7-FboxD-HF | Bsal | pICH47742, pICSL51277,<br>pICSL50001, pICH41414 | Carb |
| OsTLP7-TLD<sup>PIP2</sup>-HF | Bsal | pICH47742, pICSL51277,<br>pICSL50001, pICH41414 | Carb |
| OsTLP7-mCherry | Bsal | pICH47742, pICSL51277,<br>pICSL50004, pICH41414 | Carb |
| CFP-NLS | Bsal | pICH47742, pICSL13002,<br>pICSL30043, pAGM3205,<br>pICH41414 | Carb |
| OsTLP7-TLD-mCherry | Bsal | pICH47742, pICSL51277,<br>pICSL50004, pICH41414 | Carb |
| OsTLP7-FboxD-mCherry | Bsal | pICH47742, pICSL51277,<br>pICSL50004, pICH41414 | Carb |
| OsTLP7-TLD<sup>PIP2</sup>-mCherry | Bsal | pICH47742, pICSL51277,<br>pICSL50004, pICH41414 | Carb |
| Tie1<sup>Δ53</sup>-HA | Bsal | pICH47742, pICSL51277,<br>pICSL50009, pICH41414 | Carb |
| OsTLP7-TLD<sup>C331E</sup>-HF | Bsal | pICH47742, pICSL51277,<br>pICSL50004, pICH41414 | Carb |
| OsTLP7-TLD<sup>F336E</sup>-HF | Bsal | pICH47742, pICSL51277,<br>pICSL50004, pICH41414 | Carb |
| OsTLP7-TLD<sup>V340E</sup>-HF | Bsal | pICH47742, pICSL51277,<br>pICSL50004, pICH41414 | Carb |
| OsTLP7-TLD<sup>N347E</sup>-HF | Bsal | pICH47742, pICSL51277,<br>pICSL50004, pICH41414 | Carb |
| Tie1<sup>Y56R</sup>-HA | Bsal | pICH47742, pICSL51277,<br>pICSL50009, pICH41414 | Carb |
| Tie1<sup>V58E</sup>-HA | Bsal | pICH47742, pICSL51277,<br>pICSL50009, pICH41414 | Carb |
| Tie1<sup>N63R</sup>-HA | Bsal | pICH47742, pICSL51277,<br>pICSL50009, pICH41414 | Carb |
| Tie1<sup>Q65R</sup>-HA | Bsal | pICH47742, pICSL51277,<br>pICSL50009, pICH41414 | Carb |
| OsTLP7-TLD<sup>F336E</sup>-mCherry | Bsal | pICH47742, pICSL51277,<br>pICSL50004, pICH41414 | Carb |
| Tie1<sup>RD</sup>-HA | Bsal | pICH47742, pICSL51277,<br>pICSL50009, pICH41414 | Carb |
| Tie1<sup>CCH</sup>-HA | Bsal | pICH47742, pICSL51277,<br>pICSL50009, pICH41414 | Carb |
| GAL4-OsTLP7 | Bsal | pICSL22027, pICSL51277,<br>pICSL30054, pICH41414 | Carb |
| GAL4-OsTLP7-TLD | Bsal | pICSL22027, pICSL51277,<br>pICSL30054, pICH41414 | Carb |
| GAL4-OsTLP7-FboxD | Bsal | pICSL22027, pICSL51277,<br>pICSL30054, pICH41414 | Carb |
| GAL4-OsTLP7-TLD<sup>PIP2</sup> | Bsal | pICSL22027, pICSL51277,<br>pICSL30054, pICH41414 | Carb |
| GAL4-OsTLP7-HF | Bsal | pICSL22027, pICSL51277,<br>pICSL30054, pISCL50001,<br>pICH41414 | Carb |
| GAL4-OsTLP7-TLD-HF | Bsal | pICSL22027, pICSL51277,<br>pICSL30054, pISCL50001,<br>pICH41414 | Carb |
| GAL4-OsTLP7-FboxD-HF | Bsal | pICSL22027, pICSL51277,<br>pICSL30054, pISCL50001,<br>pICH41414 | Carb |
| GAL4-OsTLP7-TLD<sup>PIP2</sup>-HF | Bsal | pICSL22027, pICSL51277,<br>pICSL30054, pISCL50001,<br>pICH41414 | Carb |
| GFP-AVR-Pii - *A. thaliana* | Bsal | pISCL4723, pICSL13004,<br>pICSL30006, pICH77901 | Carb |
| Tie1-GFP- *A. thaliana* | Bsal | pISCL4723, pICSL51277,<br>pICSL50008, pICH41414 | Carb |
| GFP- *A. thaliana* | Bsal | pISCL4723, pICSL13004,<br>pICSL30006, pICSL50028,<br>pICH77901 | Carb |

